# Targeting adipocyte ESRRA promotes osteogenesis and vascular formation in adipocyte-rich bone marrow

**DOI:** 10.1101/2023.08.14.552932

**Authors:** Tongling Huang, Zhaocheng Lu, Zihui Wang, Lixin Cheng, Lu Gao, Jun Gao, Ning Zhang, Chang-An Geng, Xiaoli Zhao, Huaiyu Wang, Chi-Wai Wong, Kelvin W K Yeung, Haobo Pan, William Weijia Lu, Min Guan

## Abstract

Ectopic bone marrow adipocytes (BMAds) accumulation occurring under diverse pathophysiological conditions leads to bone deterioration. Estrogen-related receptor α (ESRRA) is a key regulator responding to metabolic stress. Here, we show that adipocyte-specific ESRRA deficiency rescues osteogenesis and vascular formation in adipocyte-rich bone marrow due to estrogen deficiency or obesity. Mechanistically, adipocyte ESRRA interferes with E2/ESR1 signaling resulting in transcriptional repression of secreted phosphoprotein 1 (*Spp1*); and positively modulates *Leptin* expression by binding to its promoter. ESRRA abrogation results in enhanced SPP1 and decreased LEPTIN secretion from both visceral adipocytes and BMAds, concertedly dictating bone marrow stromal stem cell fate commitment and restoring type H vessel formation, constituting a feed-forward loop for bone formation. Pharmacological inhibition of ESRRA protects obese mice against bone loss and high marrow adiposity. Thus, our findings highlight a therapeutic approach via targeting adipocyte ESRRA to preserve bone formation especially in detrimental adipocyte-rich bone milieu.

**Graphic abstract:** 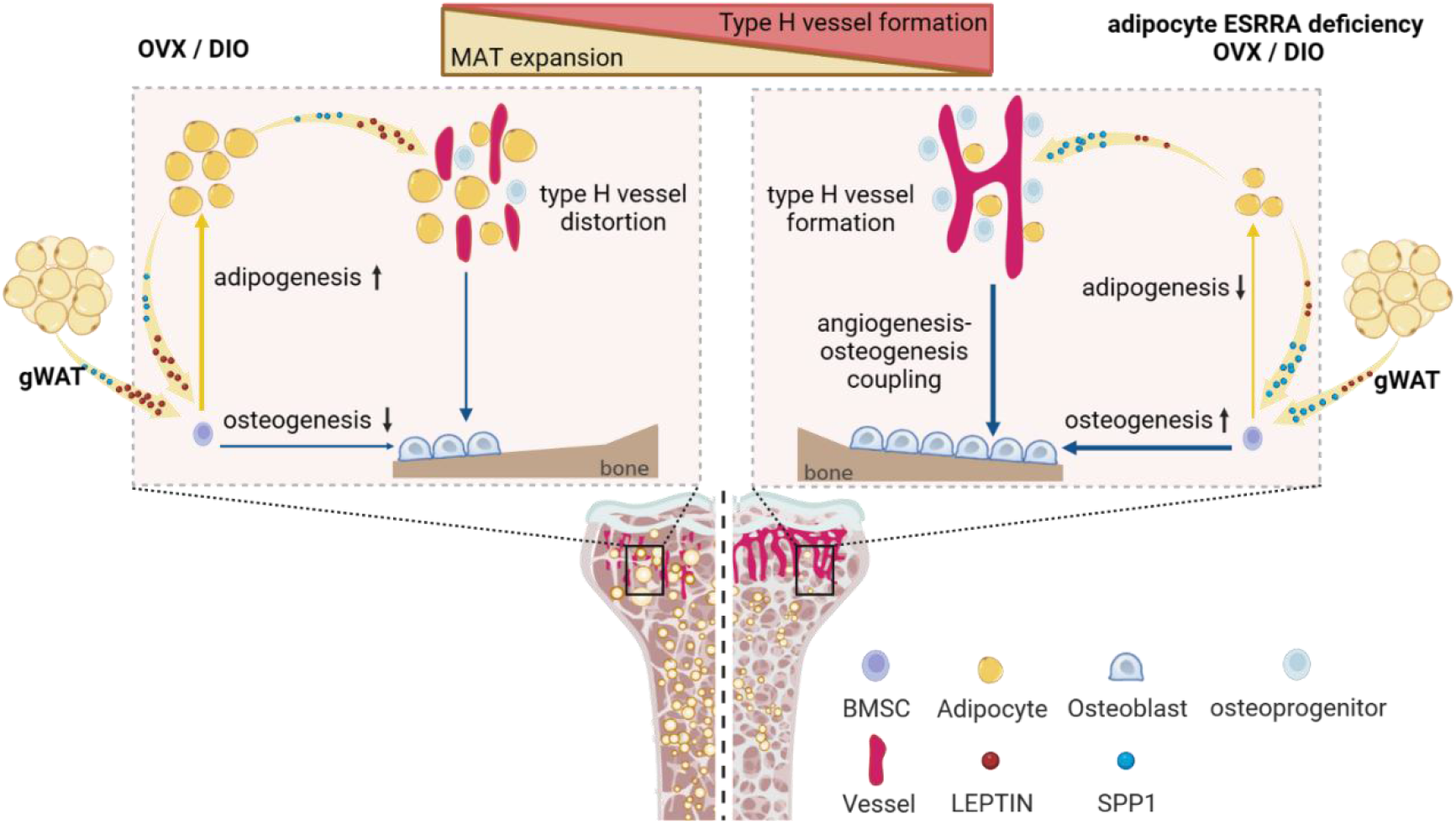

## Introduction

Mammalian bone marrow consists of multiple cell types such as adipocytes, osteoblasts, osteoclasts, stromal cells and vascular cells. The interactions of these cells provide a critical regulatory milieu for the differentiation of bone marrow skeletal (also known as stromal or mesenchymal) stem cells (BMSCs) and other lineage cells, in turn, maintaining a complex homeostatic system for remodeling and regeneration in bone microenvironment^1,2^. It is well-accepted that bone marrow adipocytes (BMAds) originate from BMSCs which also give rise to osteoblasts and osteocytes^3^. Marrow adipose tissue (MAT) accounts for approximately 10% of the total fat mass and about 70% of the whole bone marrow volume in healthy adult humans^4^. However, ectopic BMAds accumulation often occurs in diverse clinical conditions including obesity, diabetes, anorexia nervosa, glucocorticoid treatment, radiotherapy, menopause and ageing, most of which are concomitant with bone deterioration^5^. MAT expansion may exaggerate the detrimental bone microenvironment and worsen disease progression. However, the precise mechanisms of excessive BMAds particularly in response to pathophysiological conditions in mediating bone-resident cellular communication remain elusive.

Adipose tissue is central to the regulation of whole-body energy homeostasis. Similar to white adipose tissue (WAT), normal MAT exhibits energy storage function and can release FFA to support exercise-stimulated bone formation^6^. However, MAT expands rapidly in response to caloric restriction indicating that MAT is distinct from peripheral adipose tissues. Moreover, a high-fat diet also rapidly increases MAT expansion through circulating LEPTIN acting on Leptin receptor expressing (LepR^+^) BMSCs to promote adipogenesis and inhibit osteogenesis of BMSCs^3,7^. Thus, it is plausible that aberrant signaling within bone microenvironment as a result of pathophysiological conditions drives BMSC lineage allocation toward committed adipogenic progenitors at the expense of osteoprogenitors, resulting in bone deterioration. Recent studies have described that accumulated BMAds contribute to circulating signal factors such as DPP4, RANKL and several well-known adipokines such as ADIPONECTIN and LEPTIN in response to obesity, ageing or caloric restriction^2,8,9,10,11^. These secreted factors from either peripheral WAT or MAT might further impact on bone homeostasis.

BMAds are located in the bone milieu and in close contact with vascular and hematopoietic tissues. Osteogenesis is tightly coupled to bone angiogenesis; especially, a specific capillary endothelial cell (EC) subtype termed type H blood vessel in the metaphysis is linked to bone marrow vasculature at the metaphyseal-diaphyseal interface^12^. Senescence of BMAds triggered by glucocorticoid causes secondary senescence of surrounding vascular cells and osteoblasts leading to bone defect^13^. Abnormal type H vessels are also observed in aged individuals and postmenopausal women suffering from osteoporosis which is frequently accompanied by augments in marrow adiposity^14,15^. These evidences largely reflect that BMAds, osteoblasts and vessels are engaged in a reciprocal relationship maintaining bone microenvironment homeostasis. In fact, adipocyte-rich marrow is believed to impose a dominant negative effect on hematopoiesis after irradiation^16^. However, BMAds arising from Adiponectin-Cre labeled progenitors have recently been identified as the source of stem cell factor (SCF) which is required for normal haematopoiesis in young adult mice. Moreover, *Scf* transcripts are highly enriched in BMAds of human bone marrow from young donors, suggesting that BMAds play essential roles in normal marrow microenvironment^17^. Recent single cell RNA-seq based evidence further proposes that a subpopulation of Adipoq-labled marrow adipocytes is critical for supporting marrow vasculature^18^. Thus, there are context dependent functions of BMAds as a niche component maintaining bone marrow microenvironment.

Orphan nuclear receptor estrogen-related receptor α (ESRRA; also known as ERRα or NR3B1) is an essential transcription factor governing energy balance and metabolism^19^. Specifically, ESRRA is required in stress-induced response to fasting, calorie restriction, cold exposure or overnutrition^20,21,22^. We previously found that liver-specific ESRRA knockout results in decreased levels of blood lipids due to blunted hepatic triglyceride-rich very low-density lipoprotein secretion in fasting mice^23^. ESRRA is also characterized as a key mediator involved in regulating diurnal metabolic rhythm-dependent intestinal lipid absorption, coupling cell metabolism and differentiation of kidney proximal tubule or controlling liver metabolism in response to insulin action and resistance^24,25,26^. Here, we generated adipocyte-specific *Esrra* knockout mice by using an Adiponectin recombinase (Adipoq-Cre) which labels most BMAds and also peripheral adipose depots^2,11,17^. We found that adipocyte ESRRA deficiency protected mice against bone loss and high marrow adiposity induced by either a high-fat diet (HFD) or ovariectomy (OVX) with little effect on peripheral WAT. Using *in vitro* and *in vivo* approaches, we established that adipocyte ESRRA inhibition promotes bone formation and reforms bone vasculature by oppositely regulating the expression and secretion of LEPTIN and secreted phosphoprotein 1 (SPP1, also known as OSTEOPONTIN).

## Results

### Conditional adipocyte deletion of ESRRA enhances bone formation and counteracts marrow fat accumulation in DIO osteopenia mice

We utilized a conditional knockout strategy by intercrossing *Esrrα*^fl/fl^ mice with Adipoq-Cre to generate adipocyte-specific *Esrrα* knockout mice (*Esrrα*^fl/fl^; *AdipoqCre,* denoted *Esrrα*^AKO^) (Supplementary Fig. S1A, B). *Esrra*^AKO^ mice at the age of 10 weeks developed normally with no significant differences in growth, body weight, and WAT phenotype compared to *Esrrα^fl/fl^* littermates (Supplementary Fig. S1C-E). MicroCT analysis of distal femurs of *Esrra*^AKO^ male mice showed no changes in trabecular bone volume per total volume (BV/TV), trabecular bone number (Tb.N) and trabecular bone thickness (Tb.Th) compared to control littermates; in addition, no changes were detected in plasma concentrations of procollagen type 1 N-terminal propeptide (P1NP, a bone formation marker) (Supplementary Fig. S1F-H). These data suggest that *Esrrα* deletion in adipocytes did not affect the phenotypes of WAT and bone in adult mice under normal physiological condition.

Obesity is believed to induce ectopic adipocyte accumulation not only in peripheral WAT but also in bone marrow cavities in rodents and humans, contributing to detrimental disturbances in bone remodeling or bone regeneration^2,27^. Since ESRRA is responsive to stress-induced metabolic challenges including overfeeding^21,26^, we queried whether adipocyte-specific ablation of ESRRA is resistant to diet-induced obesity (DIO) or involved in fat-bone axis. Compared to *Esrrα*^fl/fl^ DIO mice, *Esrra*^AKO^ DIO male mice are similar in WAT mass and adipocyte size, and plasma levels of TG and FFA under fasting condition (Fig. 1A-F). Moreover, these two groups of mice showed similar glucose disposal rates upon an oral glucose tolerance test (Fig. 1G). On the other hand, the circulating levels of LEPTIN, which is proportional to fat stores and considered a mature adipocyte marker, were significantly reduced in *Esrra*^AKO^ DIO mice (Fig. 1H). Notably, the trabecular bone mass in *Esrra*^AKO^ DIO mice was greater than that of *Esrrα*^fl/fl^ littermates (Fig. 1I). *Esrra*^AKO^ DIO mice had significant increases in BV/TV, Tb.N and Tb.Th; whereas, Tb.Sp was reduced (Fig. 1J). In addition, *Esrra*^AKO^ DIO mice had more osteoblasts (Ob.N/BS and Ob.S/BS), indicating an enhancement of bone formation confirmed by significant increases in trabecular mineral apposition rate (MAR) and bone formation rate (BFR/BS) using calcein double labeling (Fig. 1K and 1L). However, bone histomorphometry analysis of femoral metaphyses displayed no difference in bone resorption between the two groups revealed by TRAP staining, quantification of osteoclast number (Oc.N/BS) and proportion of bone surface with active osteoclasts (Oc.S/BS) (Fig. 1M). We detected no changes in cortical bone including the midshaft cortical BV/TV and cortical bone thickness (Ct. Th) (Supplementary Fig. S2A, B). Consistently, plasma concentrations of P1NP were apparently elevated in *Esrra*^AKO^ DIO mice but collagen type I c-telopeptide (CTX-1, a bone resportion marker) remained unchanged, suggestive of increased net bone formation in *Esrra*^AKO^ DIO mice (Fig. 1N, O). As expected, HFD promoted MAT expansion. However, histological examination in femur sections by immunofluorescence staining of PERILIPIN1 (PLIN1), an adipocyte marker, and SPP1, a bone matrix protein marker, revealed that loss of ESRRA dramatically restrained HFD-induce MAT expansion while promoting bone matrix formation (Fig. 1P). Collectively, these data demonstrated that adipocyte ESRRA ablation counteracts DIO-associated bone loss and high marrow adiposity by favoring bone formation.

**Figure 1.**
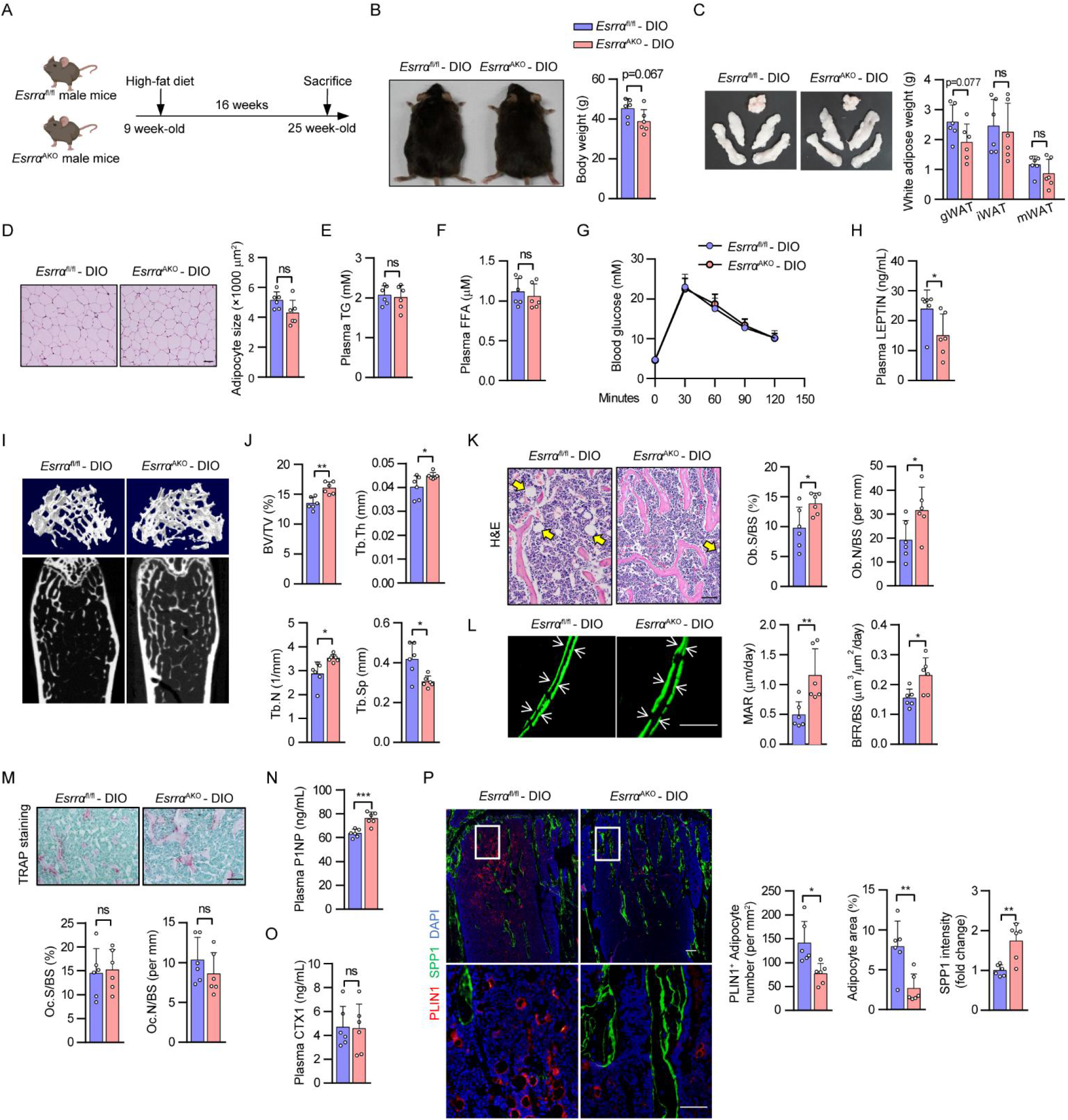
Adipocyte-specific ESRRA ablation augments bone formation and inhibits MAT expansion in DIO osteopenia mice. (A) Schematic diagram illustrating the experimental procedure for diet-induced obesity (DIO) mice model. 9-week-old *Esrra*^fl/fl^ and *Esrra*^AKO^ male mice were fed a high-fat diet (HFD) for 16 weeks. (B) Representative pictures and body weights of DIO mice. (C) Representative images and weight analysis of white adipose tissue (WAT) depots, including gonadal WAT (gWAT), inguinal WAT (iWAT), and mesentery WAT (mWAT). (D) H&E-stained gWAT sections (scale bar: 50 μm). The areas of adipocytes size are presented as graphs. (E-F) Plasma TG (E) and FFA (F) levels. (G) Oral glucose tolerance test (OGTT) was analyzed in *Esrra*^fl/fl^ and *Esrra*^AKO^ mice fed a HFD for 12 weeks. (H) Plasma LEPTIN levels. (I) Representative micro-CT images of the distal femoral trabecular bone. (J) Quantitative analysis of bone volume/tissue volume ratio (BV/TV), trabecular thickness (Tb. Th), trabecular number (Tb. N) and trabecular separation (Tb. Sp). (K) H&E staining of femur sections (scale bar: 100 μm). Yellow arrows indicate the bone marrow adipocytes. Osteoblast surface to bone surface ratio (Ob.S/BS) and osteoblast number to bone surface ratio (Ob.N/BS) are shown on the right panel. (L) Calcein double labeling of trabecular bone (scale bar: 20 μm). Mineral apposition rate (MAR) and bone formation rate (BFR/BS) were determined. White arrows indicate the distance between calcein double labeling. (M) TRAP staining of femur sections with quantitative analysis of Oc.S/BS and Oc.N/BS. TRAP-positive purple spots indicate multinucleated osteoclasts (scale bar: 100 μm). (N-O) Plasma P1NP (N) and CTX1 (O) levels. (P) PLIN1 positive marrow adipocytes (PLIN1^+^, red) and SPP1 (green) immunofluorescence staining in femur sections (scale bar: 100 μm). The box in the upper showing the metaphysis region near growth plate is represented at higher magnification in the bottom (scale bar: 50 μm). The numbers and areas of adipocytes in the femur marrow per tissue area and the quantification of SPP1 fluorescence intensity were measured. Data are shown as mean ± SD (n = 6 mice). \**P* < 0.05 was considered statistically significant.

### Loss of *Esrrα* in adipocytes attenuates bone loss through favoring bone formation and inhibiting marrow adiposity in OVX-induced osteoporosis

There is a high coincidence rate of bone loss and excessive fat deposition including MAT expansion in postmenopausal women^28,29^. Similarly, estrogen deprivation also leads to fat-bone imbalance in rodents^30^. To investigate whether lack of *Esrrα* in adipocytes affects bone homeostasis under hormonal deficiency, ovariectomized (OVX) female mice were utilized to mimic estrogen deficiency-related osteoporosis. As expected, 18-wk-old female mice at 8 weeks after OVX gained body weight with excessive WAT deposition in *Esrrα*^fl/fl^ controls (Fig. 2A-C, Supplementary Fig. 3A). However, there are no differences in body weight, WAT phenotype, blood lipids and glucose between the two genotypes in sham or OVX groups (Fig. 2B, C and Supplementary Fig. 3A-D). These evidences implied that adipocyte ESRRA abrogation did not alter WAT development upon estrogen deprivation. Similar to *Esrra*^AKO^ DIO mice, the increase in circulating LEPTIN induced by OVX was significantly reduced in *Esrra*^AKO^ OVX mice despite no differences in peripheral WAT mass (Fig. 2D).

**Figure 2.**
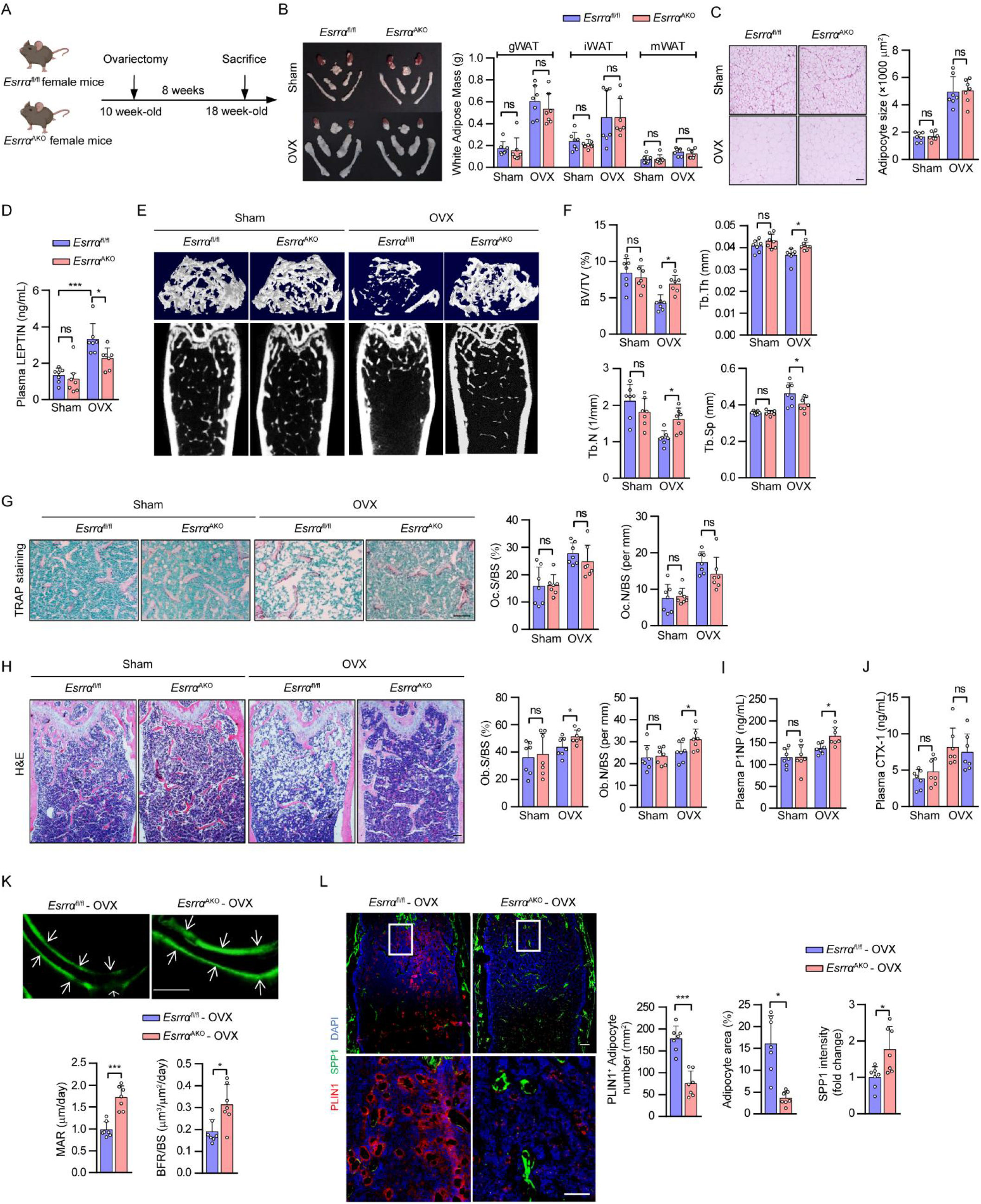
ESRRA deficiency in adipocytes favors bone formation and attenuates marrow adiposity in OVX-induced osteoporosis. (A) Schematic diagram illustrating the experimental procedure for ovariectomy (OVX)-induced osteoporosis mice model. 10-week-old *Esrra*^fl/fl^ and *Esrra*^AKO^ female mice underwent either sham or OVX operation for 8 weeks. (B) Representative images and weights of adipose depots. (C) Representative images and adipocytes size analysis from H&E-stained gWAT sections (scale bar: 50 μm). (D) Plasma LEPTIN levels. (E-F) Micro-CT images of distal femurs in sham and OVX mice (E) with morphometric analysis of BV/TV, Tb.N, Tb.Th, and Tb.Sp (F). (G) Representative TRAP-stained images and quantification of Oc.S/BS and Oc.N/BS in distal femoral metaphysis regions from sham and OVX mice (scale bar: 100 μm). (H) Representative H&E-stained images and quantification of Ob.S/BS and Ob.N/BS (scale bar: 100 μm). (I-J) Plasma P1NP (I) and CTX1 (J) levels. (K) Calcein double labeling with quantitative analysis of MAR and BFR/BS (scale bar: 20 μm). (L) Immunofluorescence co-staining and quantification of PLIN1^+^ bone marrow adipocytes (red) and SPP1 (green) of femur sections from OVX mice. Scale bar: upper panel, 100 μm; lower panel, 50 μm. Data are shown as mean ± SD (n = 7 mice). \**P* <0.05 was considered statistically significant.

MicroCT analysis of the femur trabecular bone showed that the disruption in bone remodeling induced by OVX measured by BV/TV, Tb.N, Tb.Sp and Tb.Th was significantly ameliorated in *Esrra*^AKO^ mice in comparison to *Esrrα*^fl/fl^ controls (Fig. 2E, F). No differences in bone resorption and cortical bone parameters were found between *Esrra*^AKO^ OVX mice and the corresponding controls (Fig. 2G and Supplementary Fig. 3E, F). However, osteoblast number and osteoblast surface were noticeably elevated in *Esrra*^AKO^ OVX mice due to accelerated bone formation (Fig. 2H). Consistently, blood levels of P1NP and CTX-1 further confirmed the increased net bone formation in *Esrra*^AKO^ mice following OVX (Fig. 2I, J) evidenced by increases in MAR and BFR (Fig. 2K). As expected, OVX significantly increased MAT accumulation accompanied with bone loss (Fig. 2H). Despite unchanged extramedullary WAT phenotype in *Esrra*^AKO^ OVX mice, MAT expansion induced by estrogen deprivation was substantially eliminated in mutant OVX mice (Fig. 2B, H, L). *Esrra*^AKO^ OVX mice displayed marked declines in the number and size of marrow adipocytes while having more osteoblasts around trabecular bone (Figure 2L). These evidences reveal that adipocyte-specific ESRRA deficiency has a major impact on facilitating bone formation and inhibiting marrow adipogenesis under estrogen deficiency.

### Adipocyte-specific ESRRA ablation leads to increased SPP1 expression

The fact that lack of ESRRA in adipocytes apparently reduces marrow adiposity in DIO and OVX mice drew our attention to BMAds. Given that BMAds originate from BMSCs present in the bone marrow, ESRRA expression was confirmed to be abrogated in *Esrra*^AKO^ BMAds lineage cells at indicated days after adipogenic induction (Fig. 3A). Then, isolated BMSCs from *Esrra*^fl/fl^ and *Esrra*^AKO^ mice were differentiated for 14 days in adipogenic or osteogenic induction medium, but no significant differences were observed (Fig. 3B, C). These data suggest that Adipoq-Cre-driven ablation of *Esrra* did not directly change the differentiation capability of BMSCs under normal physiological condition which is consistent with *in vivo* evidences. Treatment of rosiglitazone (Rosi) or pioglitazone can dramatically induce the expression of peroxisome proliferator activated receptor gamma (PPARG) target genes and exaggerate MAT accumulation in rodents or humans ^13,18,31^. Therefore, we induced BMSCs differentiation by Rosi to mimic adipocyte expansion under pathological conditions and observed a reduction in adipogenesis of BMSCs from *Esrra*^AKO^ mice (Fig. 3C). These evidences indicated that loss of ESRRA during adipocyte expansion under pathological conditions might exert a secondary or paracrine effect on heterogeneous BMSCs differentiation.

**Figure 3.**
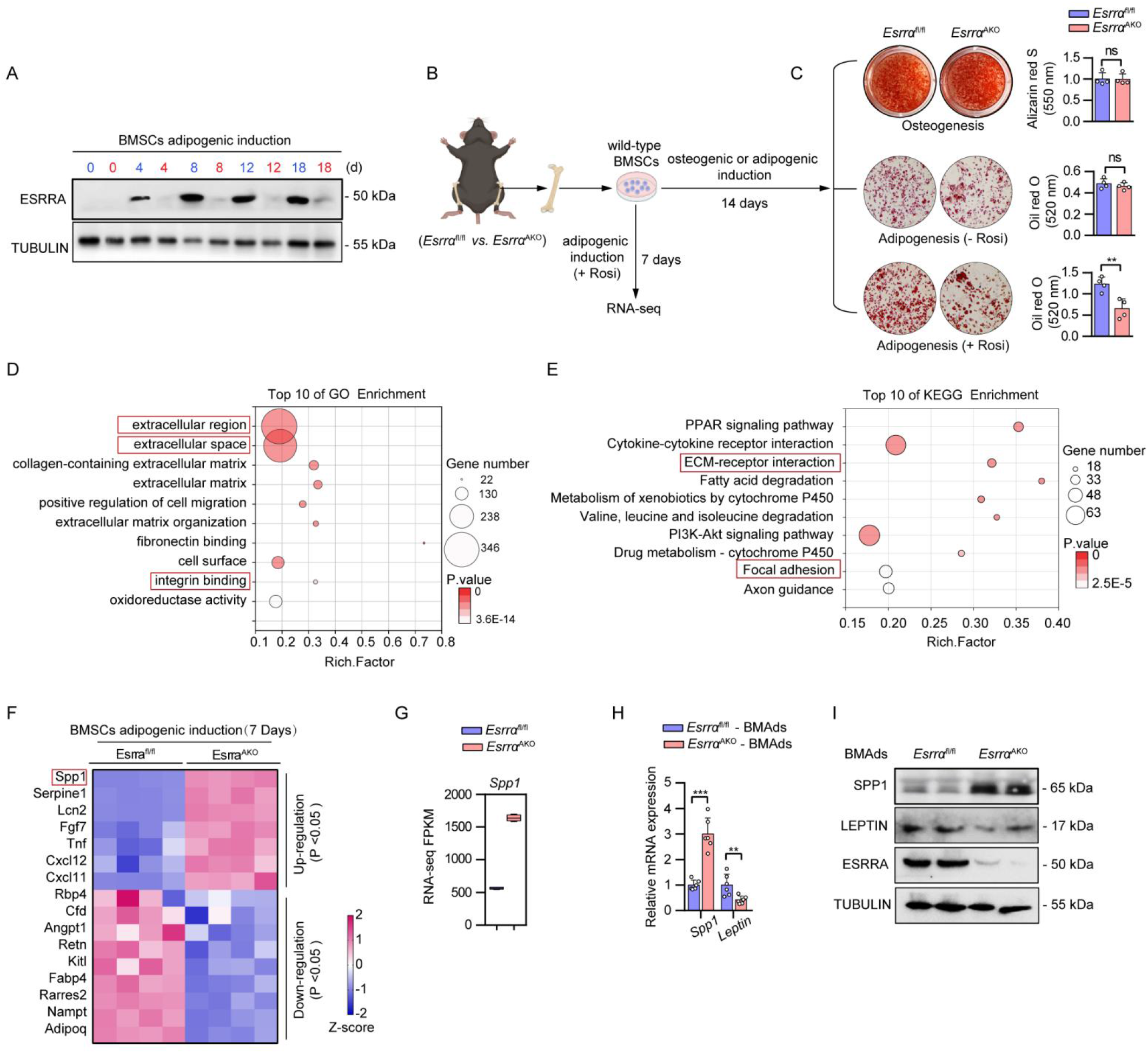
Loss of ESRRA results in enhanced SPP1 expression in adipocytes. (A) Protein expression levels of ESRRA were evaluated in BMSCs upon adipogenic induction for indicated days, comparing *Esrra*^fl/fl^ mice (blue font) with *Esrra*^AKO^ mice (red font). (B) Schematic representation of the experimental design. BMSCs were isolated from *Esrra*^fl/fl^ and *Esrra*^AKO^ mice and subsequently subjected to either adipogenic or osteogenic induction for indicated days. (C) Representative images and quantitative analyses of alizarin red S staining and oil red O staining following the indicated induction. Rosi, rosiglitazone. (D-E) Gene Ontology (GO) (D) and Kyoto Encyclopedia of Genes and Genomes (KEGG) (E) pathway enrichment analyses of all differentially expressed genes by RNA-seq (top 10 according to adjusted *P* value). (F) Heatmap depicting selected genes related to secreted factors. n = 4 per group. (G) Boxplot showing the transcript expression value (FPKM) of *Spp1* based on RNA-seq data. (H-I) Validation of SPP1 and LEPTIN expression were performed by quantitative real-time polymerase chain reaction analysis (qRT-PCR) (H) and western blotting analysis (I) in BMAds that were fully differentiated for 14 days. n = 6 per group. All the data are shown as mean ± SD. \**P* <0.05 was considered statistically significant.

To explore the transcriptional role of ESRRA in regulating BMAds autonomous genes, bulk RNA-seq transcriptome analysis was performed in BMAds lineage cells at 7 days upon adipogenic induction (Fig. 3B). By using Gene Ontology (GO) and Kyoto Encyclopedia of Genes and Genomes (KEGG) analysis, differentially expressed genes were found to be enriched in pathways related to extracellular environment and cellular response, indicating the involvement in cell signaling and interactions (Fig. 3D, E). Given that multiple secreted factors from BMAds are well-known functional regulators^32^, priority was given to 16 differentially expressed secreted factors (Fig. 3F). Notably, SPP1 was most robustly upregulated among the top-ranked secreted factors by ESRRA ablation (Fig. 3F, G). SPP1 is a well-known matrix protein by binding to multiple integrins through a highly conserved arginine-glycine-aspartic acid (RGD) motif^33^ and involved in multiple pathways including the extracellular region, integrin binding, ECM-receptor interaction, and focal adhesion (Fig. 3D, E). Furthermore, qRT-PCR profiling and western blot analysis confirmed that SPP1 expression was indeed enhanced dramatically by ESRRA abrogation in fully differentiated BMAds. In contrast, the expression levels of LEPTIN and other adipogenic marker genes such as *Pparg*, *Cebpa* and *Fabp4* were down-regulated (Fig. 3H, I and Supplementary Fig. S4). Taken together, these *ex vivo* data reveal that ESRRA ablation in BMAds led to altered expression of a panel of secreted factors, especially a significant augment of SPP1 expression.

### Adipocyte ESRRA negatively regulates the transcription of *Spp1* by interfering with E2/ESR1 signaling

We next sought to dissect the mechanism by which ESRRA negatively regulates SPP1 expression in adipocytes. Compared to sham, circulating SPP1 was significantly decreased in OVX mice but was partially rescued in *Esrra*^AKO^ OVX mice (Figure 4A). This ESRRA-mediated SPP1 production and secretion may not only arise from BMAds but also derive from peripheral WAT. We detected local SPP1 expression in gWAT with LEPTIN as a WAT-adipocyte marker by using immunofluorescence double staining. The results showed that ESRRA knockdown increased SPP1 expression in gWAT adipocytes, concomitant with decreased LEPTIN expression confirmed by qRT-PCR and western blot analysis (Fig. 4B-D). No changes were observed in other adipocyte-related genes including *Adipoq*, *Pparg*, *Cebpa* and *Fabp4* (Figure 4D). Local SPP1 expression were remarkably reduced in gWAT adipocytes of wild-type female mice 4 and 8 weeks after OVX, suggesting estrogen might be essential for adipocyte SPP1 expression (Fig. 4E, F and Supplementary Fig. S5). These results clearly revealed that in addition to BMAds, WAT-adipocyte is also a source of SPP1. Importantly, a reduction in adipocyte SPP1 expression due to estrogen deficiency can be reversed by ESRRA ablation.

**Figure 4.**
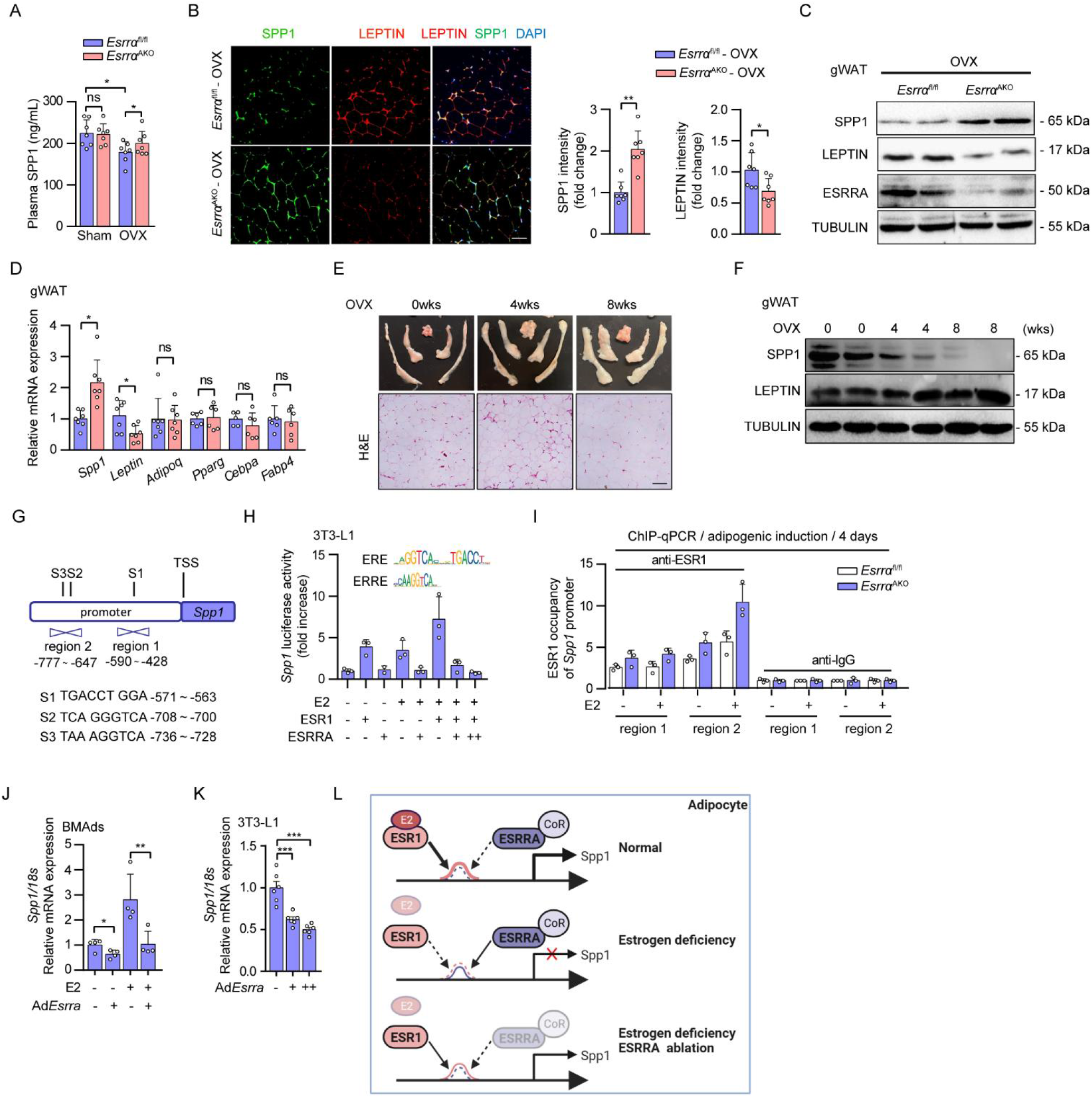
ESRRA represses SPP1 transcriptional expression via interrupting with E2/ESR1 signaling in adipocytes. (A) Plasma SPP1 levels of *Esrra*^fl/fl^ and *Esrra*^AKO^ mice at 8 weeks post-OVX or sham operation (n = 7). (B) Representative images and analysis of SPP1 and LEPTIN co-staining of gWAT from OVX mice studied in (A). (C) Protein expression analysis of SPP1, LEPTIN and ESRRA of gWAT from OVX mice. (D) mRNA expression of *Spp1*, *Leptin*, *Adipoq*, *Pparg*, *Cebpa* and *Fabp4* of gWAT from OVX mice (n = 7). (E) Representative images of white adipose depots were obtained from wild-type mice at 0 week, 4 weeks, and 8 weeks after OVX surgery. (F) Immunoblot analysis of SPP1 and LEPTIN in gWAT from the same mice studied in (E). (G) Schematic diagram displays the potential binding sites of ESR1 within the *Spp1* promoter, including S1, S2 and S3. Positions of qRT-PCR primers on the *Spp1* promoter used in ChIP are shown as region 1 and region 2. (H) Luciferase reporter activities of the *Spp1* promoter alone or in the presence of E2 in adipogenesis induced 3T3-L1 cells transfected with *Esrra* or *Esr1* expressing plasmids or control vector. n = 3 biologically independent samples. The consensus sequence binding motifs for ESR1 response element (ERE) and ESRRA response element (ERRE) are presented as indicated. (I) ChIP assay with ESR1 antibody or IgG in BMSCs from *Esrra*^fl/fl^ and *Esrra*^AKO^ mice after adipogenic induction for 4 days along with or without E2 (10 nM). n = 3 biologically independent samples. (J) mRNA expression levels of *Spp1* were measured in BMAds infected with adenovirus expressing *Esrra* or control *Gfp* in the presence of the indicated treatments of E2 for 2 days. n = 4 biologically independent samples. (K) mRNA expression levels of *Spp1* were measured in matured 3T3-L1 adipocytes infected with adenovirus expressing *Esrra* or *Gfp* for 2 days. n = 6. (L) Diagram illustrating the mechanism of ESRRA-regulated the repression of *Spp1* transcriptional expression via interfering with E2/ESR1 signaling in adipocytes. Data are shown as mean ± SD. \**P* <0.05 was considered statistically significant.

Although ESRRA does not bind 17β-estradiol (E2), ESRRA can modulate estrogen receptor alpha (ESR1)-mediated response of shared target genes in an E2 dependent way^34^. *Spp1* promoter does not contain any canonical ESR1 response element (ERE) with an invert repeat of AGGTCA half-sites. Nonetheless, E2/ESR1 can regulate a single half-site preceded by a trinucleotide sequence TCA, which is also an ESRRA response element (ERRE)-like sequence^35^. Three such sites (S1-S3) can be found on the *Spp1* promoter (Fig. 4G). We found that ESR1 induced *Spp1* promoter activity, which was further enhanced by E2, in 3T3-L1 preadipocytes upon adipogenic differentiation for 2 days (Fig. 4H). In contrast, ESRRA overexpression suppressed the E2/ESR1-driven *SPP1* promoter activation in a dose dependent manner, demonstrating a role of ESRRA in attenuating the E2/ESR1 dependent transcriptional activation (Fig. 4H). To further evaluate this interference, we next determined whether ESRRA knockdown would result in an induction of E2/ESR1 transactivation on the *SPP1* promoter. By chromatin immunoprecipitation (ChIP) assay, ESRRA abrogation led to enhanced recruitment of ESR1 to the *SPP1* promoter, especially region 2 which was more responsive to E2 stimulation (Fig. 4I). Consistently, mRNA expression of SPP1 was induced by E2 but was suppressed by an adenovirus expressing ESRRA in fully differentiated BMAds (Fig. 4J). Similar evidences were recaptured in 3T3-L1 mature adipocytes as well as in human BMSCs-derived BMAds (Fig. 4K, and Supplementary Fig. S6). Collectively, our data demonstrated that ESRRA negatively modulates SPP1 expression through interfering with E2/ESR1 signaling in adipocytes (Fig. 4L).

### Adipocyte ESRRA ablation results in an increase in SPP1 secretion facilitating type H vessel formation in osteoporotic mice

SPP1 is well known to exert a proangiogenic effect mediating neovascularization in the early stage of bone healing^36,37^. Notably, global knockout of SPP1 causes a reduction in type H vessels in mice long bone^38^. Therefore, it is possible that soluble form of SPP1 released from adipocytes might contribute to type H vessel formation. We performed CD31 and ENDOMUCIN (EMCN) double immunofluorescence staining in the metaphysis of distal femur to verify type H blood vessel formation. Consistent with previous studies^14,39^, OVX caused a significant reduction of type H blood vessels and adjacent Osteix^+^ osteoprogenitor cells in *Esrrα*^fl/fl^ mice compared to sham (Fig, 5A, D). In contrast, the abundance of type H vessels and osteoprogenitors were markedly preserved in *Esrra*^AKO^ mice following OVX (Fig. 5A, D). As a characteristic feature of metaphyseal type H vessels, the organization and length of columnar vessels exhibited pronounced improvement in mutant mice (Fig. 5A, B). Similarly, *Esrrα*^fl/fl^ DIO mice exhibited apparently distorted type H vessels; however, this distortion was minimal in *Esrra*^AKO^ DIO mice (Supplementary Fig. S7A, B). These results suggest that adipocyte ESRRA ablation rescues deteriorated type H vessels under pathological osteoporotic conditions while promoting perivascular osteoprogenitors accretion.

**Figure 5.**
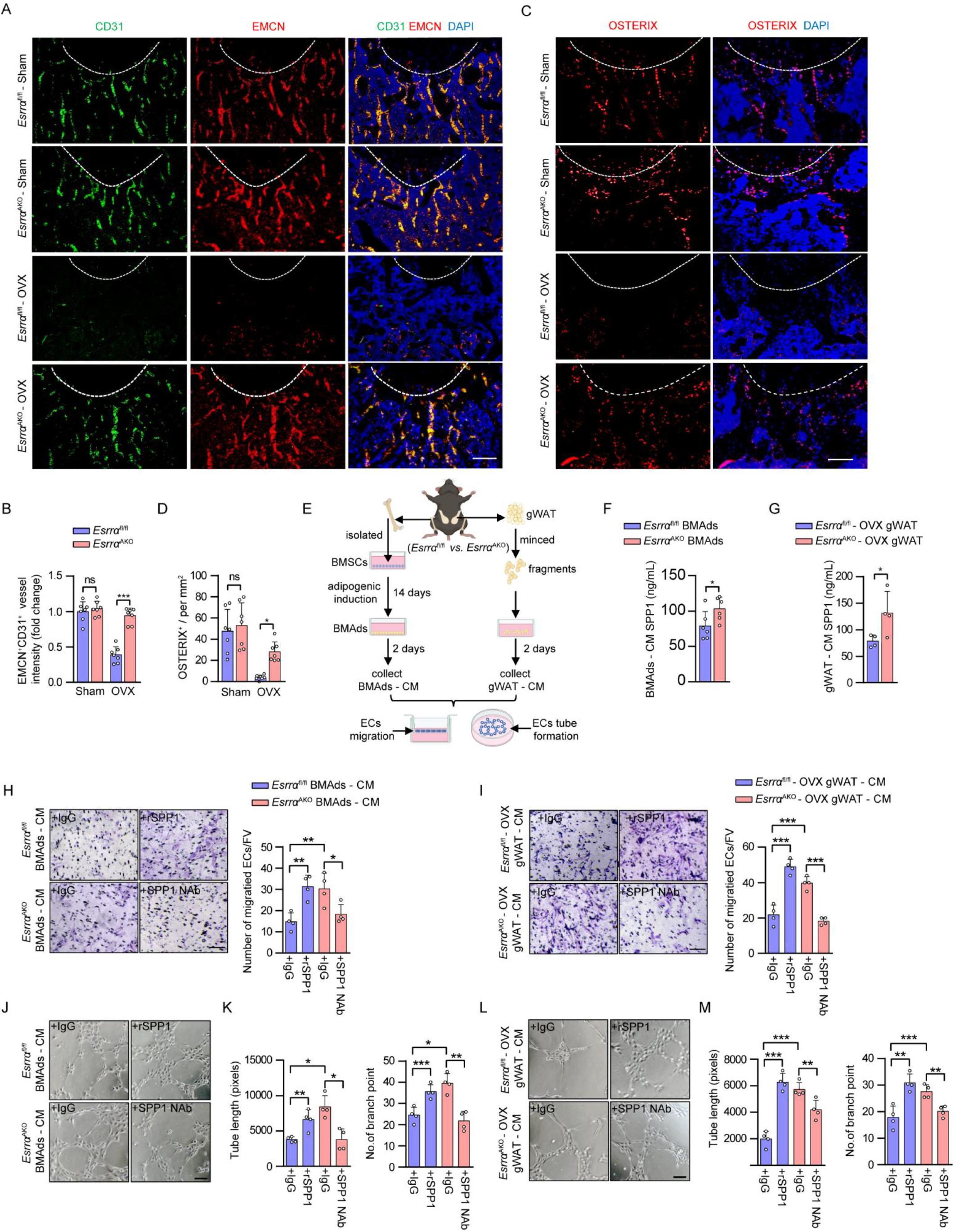
Adipocyte ESRRA deficiency rescues type H vessel formation under osteoporotic condition by facilitating SPP1-induced angiogenesis. (A) Representative images of metaphyseal type H vessels near growth plate immunostained for Endomucin (EMCN, red) and CD31 (green) in distal femurs of *Esrra*^fl/fl^ and *Esrra*^AKO^ mice following sham and OVX. DAPI (blue) is used for counterstaining of nuclei. Scale bar: 100 μm. (B) Quantification of CD31^+^ EMCN^+^ type H vessel intensity per mm^2^ (n= 7). (C) Immunostaining of OSTERIX (red) with DAPI (blue) in the metaphysis of distal femurs of *Esrra*^fl/fl^ and *Esrra*^AKO^ mice following sham and OVX. Scale bar: 50 μm. (D) Quantification of OSTERIX^+^ cells in bone marrow per mm^2^ (n = 7). (E) Schematic diagram showing the procedure of the conditioned medium (CM) preparation, tube formation assay and cell migration assay. (F-G) The concentrations of soluble SPP1 in BMAds-CM (F) or gWAT-CM (G) prepared from *Esrra*^fl/fl^ and *Esrra*^AKO^ OVX mice were measured by ELISA. n = 6 and 4, respectively. (H-I) Microvascular endothelial cell (ECs) migration in response to BMAds-CM (H) or gWAT-CM (I) with the addition of 0.5 μg/mL recombinant SPP1 (rSPP1), 1 μg/mL neutralizing SPP1 antibody (SPP1 Nab), or an equal volume of IgG for 24 hours was followed by quantification and presentation of representative images from four independent experiments (scale bars: 100 μm). (J-M) Matrigel tube formation assay was performed using ECs and BMAds-CM (J) or gWAT-CM (L) with the indicated treatments for 4 hours (scale bar: 100 μm). The length of the tubes and the number of branch points (K, M) per field were analyzed (n = 4). Data are shown as mean ± SD. \**P* <0.05 was considered statistically significant.

As a negatively charged non-collagenous matrix glycoprotein, soluble SPP1 has a high affinity for calcium allowing it to anchor in bone environment^40^. To determine whether BMAd- or WAT-derived soluble SPP1 contributed to the formation of type H blood vessels, we collected conditioned medium (CM) from both sources and performed co-culture assays to evaluate the efficacy of secreted SPP1 in inducting angiogenesis of microvascular endothelial cells (ECs) (Fig. 5E). We first detected the concentrations of SPP1 from CM and confirmed that adipocyte ESRRA abrogation enhanced SPP1 secretion (Fig. 5F, G). Transwell migration assay revealed that ECs cultured in either BMAds CM or gWAT CM from *Esrra*^AKO^ mice displayed an enhanced ability to migrate relative to *Esrrα*^fl/fl^ controls (Fig. 5H, I). Additionally, Matrigel tube formation assay confirmed that CM from both sources of mutant mice promoted the formation of capillary-like network structures (Fig. 5J, M). Importantly, recombinant SPP1 (rSPP1) rescued ECs migration and tube formation in two *Esrrα*^fl/fl^ CM groups (Fig. 5H-M). On the contrary, addition of anti-SPP1 neutralizing antibody (SPP1 NAb) in two *Esrra*^AKO^ CM groups hampered the angiogenic processes (Fig. 5H-M). These data indicated that soluble SPP1 released by both sources of ESRRA-deficient adipocytes was responsible for ECs migration and vascular formation. Taken together, our findings illustrated a potent protective effect of adipocyte ESRRA ablation on bone angiogenesis in OVX and DIO mice through elevated soluble SPP1 production. It is thus plausible that enhanced bone formation in *Esrra*^AKO^ mice is in part facilitated by the sustained bone marrow vascularization, especially under osteoporotic condition induced by estrogen derivation or overfeeding.

### Adipocyte ESRRA positively regulates transcriptional expression of *Leptin* by binding to the *Leptin* promoter

Compared to DIO or OVX controls, ESRRA deficient mice displayed declined levels of circulating LEPTIN as well as repressed expression in both WAT-adipocytes and BMAds (Fig. 1H, 2D, 3H, 4C and Supplementary Fig. S8), suggesting that *Lepin* may be a target gene of ESRRA. Thus, we evaluated whether ESRRA may directly regulate the expression of *Leptin* by binding to four putative ERREs S1-S4 on the promoter (Fig. 6A). We first established that this wild-type *Leptin* promoter is sufficient to confer, in a dose dependent manner, responsiveness to ESRRA and its co-activator PPARGC1A (Fig. 6B). Moreover, the regulation on *Leptin* promoter was dose-dependently suppressed by a well-studied ESRRA specific antagonist Compound 29 (C29) (Fig. 6C). Importantly, deletion and mutation analysis of these putative ERREs revealed that S1 and S3 ERREs were primarily responsible for the ESRRA-driven expression of *Leptin* (Fig. 6D). We next investigated if ESRRA is physically bound to these ERREs by ChIP assays using primers encompassing regions 1-3 (Fig. 6A). 3T3-L1 underwent adipogenic differentiation for 4 days to induce endogenous ESRRA and LEPTIN expression. ChIP-qPCR analysis confirmed the binding of endogenous ESRRA to regions 1 and 2 that contains S1 and S3 respectively; however, C29 reduced said ESRRA occupancy (Fig. 6E). Furthermore, augmented ESRRA occupancy was measured in regions 1 and 2 in 3T3-L1 overexpressing ESRRA (Fig. 6F). In line with these observations, LEPTIN expression was increased in a dose-dependent manner by an adenovirus overexpressing ESRRA in matured 3T3-L1 adipocyte as well as in BMAds differentiated from murine or human BMSCs (Fig. 6G, H and Supplementary Fig. S9). In summary, our data strongly established that *Leptin* is a bona fide ESRRA target gene in adipocytes.

**Figure 6.**
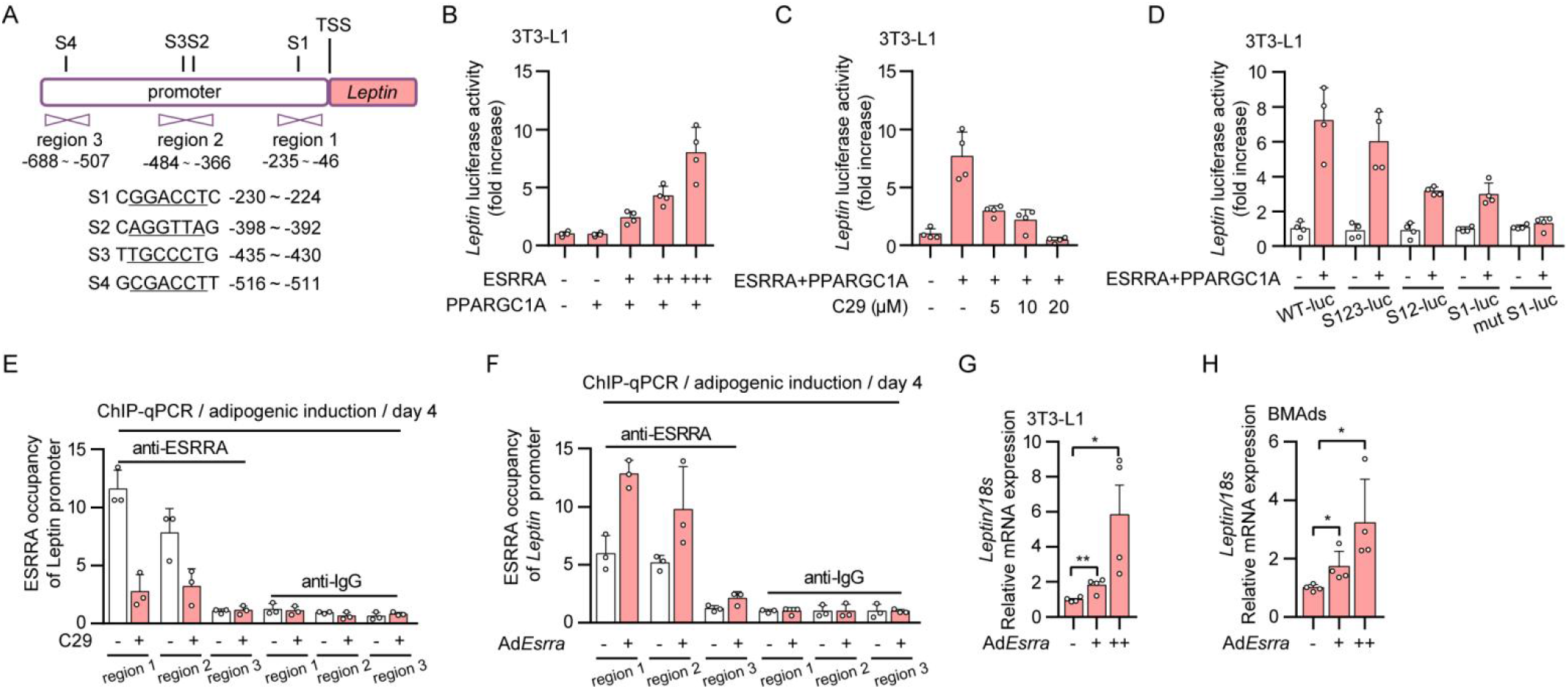
Adipocyte ESRRA positively regulates *Leptin* transcriptional expression by binding to the *Leptin* promoter. (A) Schematic diagram illustrating the putative binding sites of the ERRE on the mouse *Leptin* promoter, with four ERREs sequences highlighted by underlined nucleotides (S1, S2, S3 and S4). Positions of qRT-PCR primers on the *Leptin* promoter used in ChIP are shown as region 1, region 2 and region 3. (B) Luciferase reporter activities of the *Leptin* promoter in 3T3-L1 transiently transfected with *Esrra* and co-activator *Ppargc1a* plasmids or control vector. n = 4 biologically independent samples. (C) Effects of compound 29 (C29) on ESRRA and PPARGC1A in the regulation of the *Leptin* promoter by using luciferase reporter assays. 3T3-L1 cells were transfected with *Esrra* and *Ppargc1a* expressing plasmids and treated with the indicated doses of C29 for 24 hours. n = 4. (D) Effects of ESRRA on different constructs of *Leptin* promoter activities were tested in 3T3-L1 cells transfected with WT-luc, S123-luc, S12-luc, S1-luc, or mut S1-luc. n = 4. (E-F) ChIP assay with ESRRA antibody or IgG in 3T3-L1 cells after adipogenic induction for 4 days along with treatment of 10 μM C29 or DMSO (E), or with an infection of adenovirus expressing *Esrra* or control *Gfp* (F). n = 3. (G-H) mRNA expression of *Leptin* in matured 3T3-L1 adipocytes (G) and BMAds (H). After 14 days of adipogenic induction, matured 3T3-L1 adipocytes or BMAds were infected with adenovirus expressing *Esrra* or control *Gfp* for another 2 days. n = 4. All data in this figure are represented as mean ± SD. \**P* <0.05 was considered statistically significant.

### ESRRA ablation drives a reduction of LEPTIN and an increase of SPP1 released from adipocytes synergistically directing BMSCs lineage commitment

LEPTIN released from adipocytes acts on LepR^+^ BMSCs dictating fate commitment towards adipogenesis^3,7^. On the other hand, SPP1 directs osteogenic differentiation of BMSCs mainly in an extracellular manner via binding to integrin β1^33^. Altered BMSCs lineage commitment maybe behind the reduced BMAds and enhanced bone formation seen in *Esrra*^AKO^ mice. To assess the effects of secreted factors from distinct adipocytes on MSCs fate determination, we first collected CM from either BMAds or minced gWAT depots isolated from *Esrrα*^fl/fl^ and *Esrra*^AKO^ OVX mice (Fig. 7A). BMSCs from wild-type adult mice were then cultured in CM supplemented with osteogenic/adipogenic mixed induction medium as reported^41^ (Fig. 7A). The concentration of LEPTIN was dramatically lower in the gWAT-derived CM from *Esrra*^AKO^ OVX mice (Fig. 7B); whereas, SPP1 level was higher (Fig. 5G). In wild-type BMSCs treated with said gWAT-CM, mRNA levels of well-established osteogenic markers, *Runx2*, *Sp7* and *Bglap* were significantly up-regulated, revealing a shift toward committed osteoblast progenitors (Fig. 7C). On the contrary, adipogenic markers *Pparg*, *Cebpa* and *Fabp4* expression were significantly down-regulated, indicating a reduction in adipogenic differentiation (Fig. 7C). As expected, BMSCs treated with gWAT-CM from *Esrra*^AKO^ OVX mice exhibited a profound increase in osteogenic capability in contrast to a significant decrease in adipogenic potential (Fig. 7D, E). Compared to IgG control, when induction medium containing gWAT-CM from *Esrrα*^fl/fl^ OVX mice was supplemented with Allo-aca, a specific LEPTIN receptor antagonist peptide that blocks LEPTIN/LepR signaling, BMSCs preferentially differentiated into osteoblasts at the expense of adipocytes differentiation (Fig. 7D, E). Additionally, rSPP1 facilitated osteogenesis and inhibited adipogenesis, which was even more profound in combination with Allo-aca, demonstrating their concerted effects (Fig. 7D, E). Conversely, when recombinant LEPTIN (rLEPTIN) or SPP1 NAb was added to gWAT-CM from *Esrra*^AKO^ OVX mice, osteogenic differentiation of BMSCs was reduced; whereas, adipogenic differentiation was enhanced (Fig. 7D, E). Similar results were obtained using BMAds-CM, suggesting the presence of a paracrine effect of BMAds on surrounding BMSCs (Fig. 7F-I). Collectively, these results delineated that the loss of ESRRA in adipocytes alters the production and secretion of LEPTIN and SPP1, acting on BMSCs through either a paracrine or endocrine manner, promoting osteogenic differentiation at the expense of adipogenic differentiation; thereby, shifting the lineage commitment of BMSCs.

**Figure 7.**
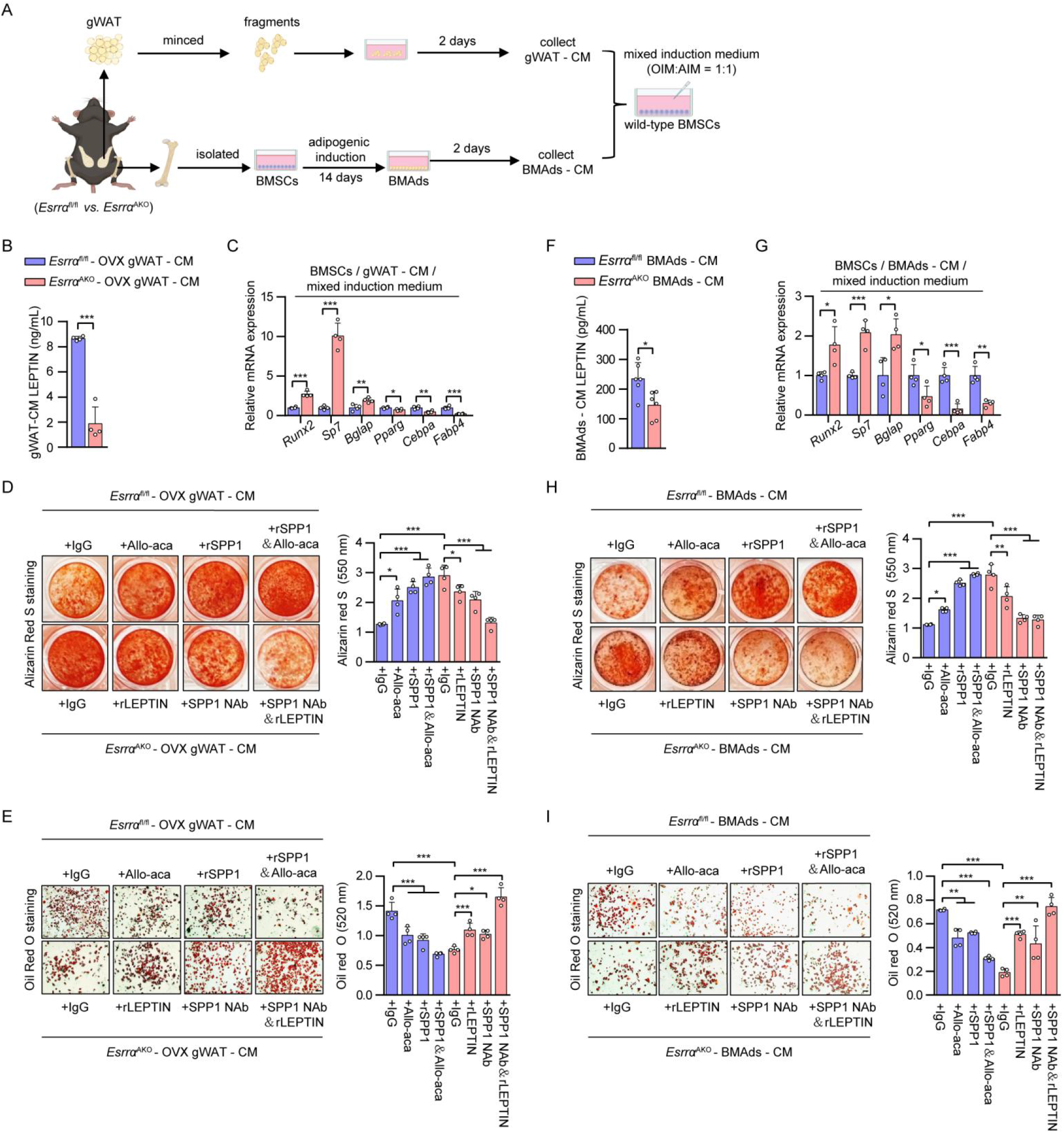
ESRRA-ablated adipocytes secrete increased SPP1 and decreased LEPTIN synergistically dictating BMSCs lineage commitment towards osteogenesis. A) Schematic diagram showing the procedure of the conditioned medium (CM) preparation from cultured BMAds or minced gWAT; and wild-type BMSCs were differentiated in osteogenic/adipogenic mixed induction medium (OIM:AIM=1:1) supplemented with the indicated CM. (B) The concentrations of soluble LEPTIN in gWAT-CM prepared from *Esrra*^fl/fl^ and *Esrra*^AKO^ OVX mice were measured by ELISA (n=4). (C) mRNA levels of osteogenesis markers *Runx2*, *Sp7*, *Bglap*, as well as adipogenic markers *Pparg*, *Cebpa*, *Fabp4* in wild-type BMSCs cultured with mixed induction medium and indicated gWAT-CM for 14 days. (D-E) Representative images and quantification of alizarin red S staining (D) and oil red O staining (E) of BMSCs cultured as in (C) with an addition of gWAT-CM for 14 days, in the presence of rSPP1, SPP1 Nab, recombinant LEPTIN (rLEPTIN), LEPTIN receptor antagonist Allo-aca or IgG as indicated. Scale bar: 100 μm. (F) The concentrations of soluble LEPTIN in BMAds-CM as prepared from (A) (n=6). (G) mRNA levels of indicated genes in wild-type BMSCs cultured in mixed induction medium supplemented with the indicated BMAds-CM for 14 days (n=4). (H-I) Representative images and quantification of alizarin red S staining (H) and oil red O staining (I) of BMSCs cultured as in (G) with the indicated treatments for 14 days (scale bar: 100 μm). The experiments were conducted according to the procedure shown in (A-E). Data are shown as mean ± SD. \**P* <0.05 was considered statistically significant.

### Pharmacological ESRRA inhibition protects bone loss and impedes MAT expansion in DIO mice

Next, we employed a specific ESRRA antagonist C29 to assess the pharmacological effect of ESRRA inhibition on adipocytes (Fig. 8A). In mature 3T3-L1 adipocytes, C29 dose-dependently suppressed LEPTIN but induced SPP1 expression while preserving adipogenesis shown by Oil Red O staining (Fig. 8B, C). Moreover, we recaptured similar results in C29-treated BMAds differentiated from murine or human BMSCs (Fig. 8D, E and Supplementary Fig. S10A); namely, C29 dose-dependently altered LEPITN and SPP1 production in an opposite manner (Supplementary Fig. S10B-E). These evidences confirmed the expression of LEPTIN and SPP1 was dictated by ESRRA but not fate switching. These data proved that *in vitro* C29 treatment can efficiently inhibit ESRRA activity in adipocytes, raising the prospect of achieving augmented therapeutic efficacy by simultaneously targeting SPP1 and LEPTIN pathways.

**Figure 8.**
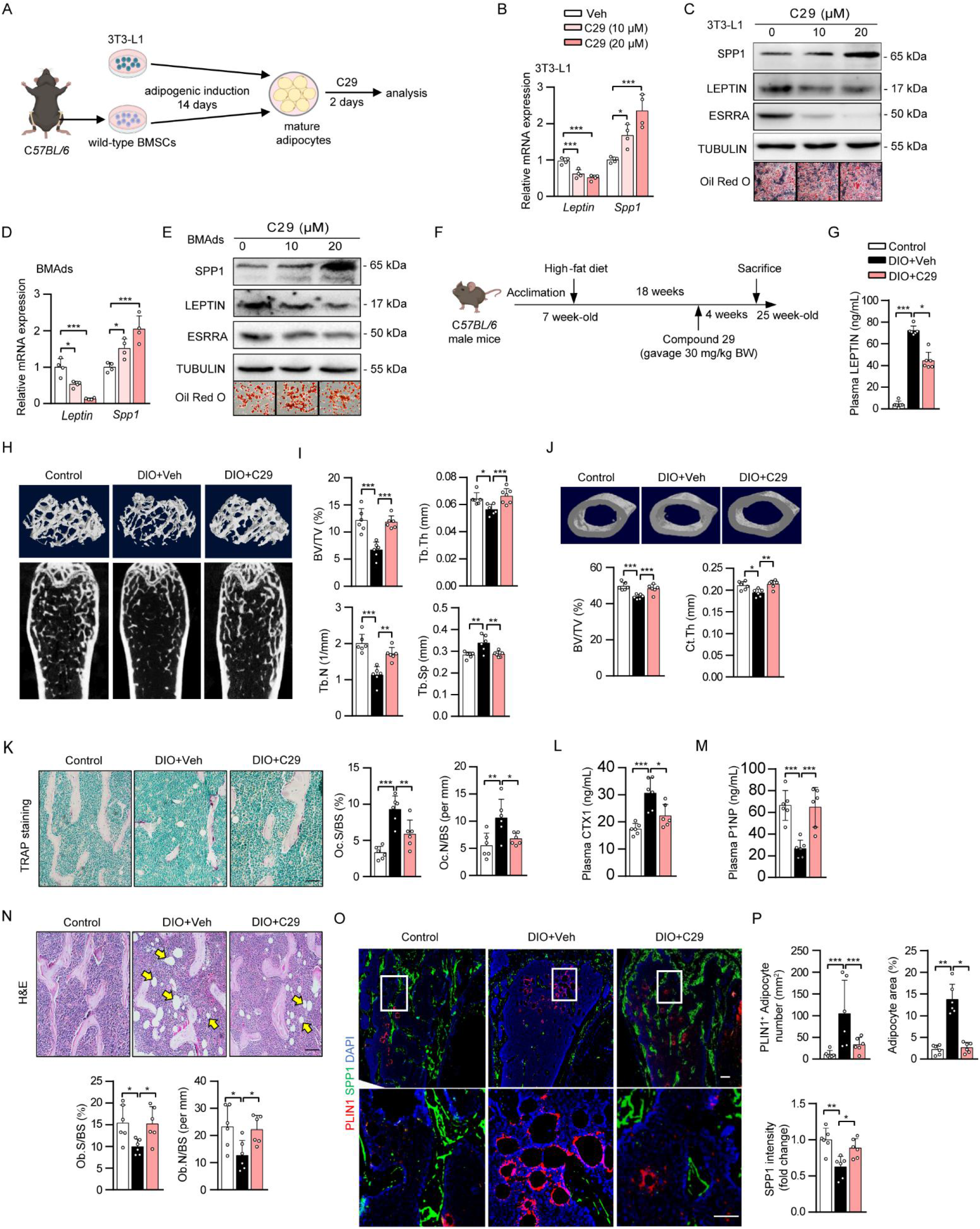
Pharmacological inhibition of ESRRA protects DIO mice from bone loss and MAT expansion. (A) Experimental design. Wild-type BMSCs were isolated from C57BL/6 mice and differentiated into BMAds, and 3T3-L1 preadipocytes were cultured in adipogenic medium for 14 days. Mature adipocytes were subsequently treated with C29 for an additional 2 days. (B-E) The mRNA and protein levels of LEPTIN and SPP1 in mature 3T3-L1 adipocytes (B-C) or BMAds (D-E). *In vitro* experiments were repeated 4 times. (F) Schematic diagram showing the experimental design for pharmacological treatments in DIO mice. Seven-week-old C57BL/6 mice were fed either a standard chow diet or HFD for 18 weeks, and received either vehicle or C29 (30 mg/kg/body weight) every day during the last 4 weeks (n = 6 mice). (G) Plasma LEPTIN levels. (H-I) Representative micro-CT images and histomorphometry analysis of BV/TV, Tb.N, Tb.Th and Tb.Sp at the distal femurs. (J) Representative micro-CT images of middle-segment of cortical bone and histomorphometry analysis of cortical bone volume/tissue volume ratio (BV/TV) and cortical thickness (Ct.Th). (K) Representative images of TRAP-stained femoral sections (scale bar: 100 μm). Quantitative assessment of trabecular Oc.S/BS and Oc.N/BS based on TRAP-stained sections (n = 6). (L-M) Plasma CTX-1 (L) and P1NP (M) levels. (N) Representative images of H&E-stained femur sections (scale bar: 100 μm). Quantitative assessment of trabecular Ob.S/BS and Ob.N/BS based on H&E-stained sections. (O) Representative PLIN1 and SPP1 immunostaining in femoral sections. Scale bar: upper panel, 100 μm; lower panel, 50 μm. (P) Quantification of PLIN1^+^ adipocyte number and of SPP1 fluorescence intensity. Data are shown as mean ± SD. \**P* <0.05 was considered statistically significant.

To further assess the pharmacological effects of C29 *in viv*o, we treated DIO male mice with C29 or vehicle for 4 weeks (Figure 8F). Consistent with our previous report^23^, C29 did not alter body weight and visceral fat mass. However, circulating LEPTIN level was markedly decreased in C29-treated compared to vehicle DIO mice (Fig. 8G). Of note, DIO mice had significant bone loss evident from μCT analysis and quantitative bone parameters which was rescued by C29 (Fig. 8H, I). Consistent with a recent study^27^, HFD decreased cortical thickness by 7% and BV/TV by 10% both were restored by C29 (Fig. 8J). This protective effect may be in part due to decreased bone resorption shown by a reduction of TRAP staining and blood CTX-1 concentration (Fig. 8K, L). This result is also consistent with ours and other previous reports that pharmacological inhibition of ESRRA directly blocks osteoclastogenesis and bone resorption^42,43^. Moreover, SPP1 immunofluorescence staining and plasma P1NP level revealed more osteoblasts and bone formation in C29-treated than vehicle DIO mice (Fig. 8M, N). Remarkably, BMAT expansion in response to HFD was diminished by C29 (Fig. 8O, P). These findings illustrate a potential therapeutic option by inhibiting ESRRA function in adipocytes for treating osteopenia associated with high marrow adiposity.

## Discussion

Bone is an active metabolic organ within which different cell types communicate with each other fine-tuning homeostasis. Marrow adipocytes enable bone-forming cells to harness essential metabolic fuels but also render them susceptible to microenvironment damages in bone pathologic niche. Here, we show that adipocyte-specific ESRRA knockout rescues bone loss and type H vessel distortion as well as inhibits marrow adiposity in mice induced by OVX or DIO. Our present analysis identifies a novel function of ESRRA in adipocytes, namely, ESRRA oppositely regulates the expression of LEPTIN and SPP1. Our findings demonstrated that blockade of ESRRA in adipocytes substantially restricted MAT expansion and promoted bone formation by altering the expression of secreted factors such as LEPTIN, SPP1 and possibly other factors which play synergistic roles on BMSCs fate determination and vascular endothelial cells angiogenesis in an adipocyte-rich bone milieu. Our results point to new avenues for treating bone disorders especially in clinical conditions associated with high marrow adiposity by chemically targeting ESRRA.

Unlike peripheral WAT and BAT, the pathophysiological role of MAT is largely unexplored. Patients with congenital generalized lipodystrophy (CGL) who lack MAT such as CGL1 and CGL2 develop pathological osteosclerosis, while patients with CGL3 and CGL4 who retain MAT fail to develop high bone density, suggesting that MAT is necessary for bone homeostasis^44^. Distinct from peripheral adipocytes, MAT can change not only size but also BMAds number in response to nutritional cues and hormonal signals. As a precursor reservoir for BMAds, MSCs are highly enriched in the bone marrow and undergo a dynamic differentiation process throughout life^3,31^. Of note, LEPIN plays an important role in regulating adipogenesis and osteogenesis through acting on LepR^+^ MSCs in the bone milieu^7^. MAT can reach approximately 30% of total fat mass under pathological conditions^8,45^ and may also contribute to LEPTIN contents. However, the endocrine or paracrine effect of MAT-derived LEPTIN is underestimated and most studies on energy balance only focus on LEPIN produced from peritoneal WAT. Noticeably, high expression of LEPIN by human bone marrow adipocytes has been observed in primary culture and that human femur MAT adipocytes secrete high levels of LEPTIN^46,47^. Importantly, LepR^+^ BMSCs were shown to be more abundant in obese human subjects associated with an enrichment of the molecular signature of adipocyte progenitor cells^48^. These observations implicate BMAds-derived LEPTIN may play a paracrine function on surrounding cells such as LepR^+^ stromal cells. In our study, we identified that LEPTIN released from BMAds imposed similar effects on directing BMSCs fate determination as LEPTIN derived from peritoneal gWAT. Our findings reinforce the negative influence of excess LEPTIN on bone homeostasis in a paracrine manner.

Triggered by excessive LEPTIN/LepR signaling, BMAds expansion may further inflate at the expense of osteoblasts since they arise from common progenitors. Indeed, our data show that circulating level of LEPIN was significantly enhanced in mice upon estrogen deficiency or overfeeding accompanied with expanded adipocytes and diminished Osterix^+^ osteoprogenitors which were rescued by ESRRA ablation in adipocytes. But peripheral WAT pads changed little by ESRRA inhibition indicating that BMSCs lineage allocation, as a highly dynamic process, is more responsive to local LEPTIN signaling in the bone microenvironment. Although we have technical difficulties in deciphering the respective contribution of LEPTIN from distinct sources, our findings from *ex vivo* adipogenic differentiation culture and conditioned medium assays strongly suggest that ESRRA inhibition in mature BMAds performed essential roles by reducing LEPIN levels and its local signaling in the bone marrow. Consistent with this concept, recently two independent studies reported that an increased local LEPTIN/LepR response driven by expanded hypodermal adipocytes leads to a chronic wound; and augmented LEPITN level in malignant tumor microenvironment facilitates benign-to-malignant transition through acting on LepR-expressing cancer stem cells^49,50^. Thus, targeting adipocyte ESRRA may also have implications to restricting the LEPTIN/LepR signaling pathway required for certain pathological progression.

In the skeletal system, vasculature is not only a conduit system for nutrients and oxygen, but also fundamental for bone formation and haematopoiesis^12,39^. Abundance of type H vessels is also considered as an important indicator of osteopenia in human subjects^14,15^. In line with previous findings, we observed that OVX mice have declined type H vessels and diminished perivascular Osterix^+^ progenitors. Type H vasculature in DIO mice was also apparently distorted and sparse. The functional properties of bone endothelial cells require cell-matrix signaling interactions frequently mediated by endothelial integrin β1 and its binding partners such as matrix glycoprotein SPP1. Loss of endothelial integrin β1 or global *Spp1* knockout results in a disorganized vasculature, that is, shorter type H vessel columns, highly branched metaphyseal capillaries and lack of a straight columnar organization^38^. We noticed that the length and organization of type H vessel columns were significantly improved by adipocyte-specific ESRRA abrogation, accompanied with a notable reduction of BMAds number and size around type H vessels below the growth plate. We revealed that SPP1 can be produced by BMAds in which *Spp1* is downregulated during adipocyte expansion, especially upon estrogen deprivation. However, ESRRA interferes with E2/ESR1-dependent regulation of *Spp1* expression as a result of their recognition of common binding sites. The repression activity of ESRRA on *Spp1* promoter might be attributed to corepressor receptor-interacting protein 140 (RIP140) recruited by ESRRA in adipocytes where RIP140 is highly expressed^51^. Thus, SPP1 production from ESRRA-deficient adipocytes may account for bone vasculature reconstruction.

In addition to its pro-angiogenic effect, SPP1 has also been attributed multiple functional roles in promoting MSC migration and adhesion, and facilitating osteogenic differentiation while retarding adipogenic differentiation through its RGD-integrin binding sequence in the soluble form^33^. Recently, a BMSC-secreted factor KIAA1199 has been identified and KIAA1199 blockade facilitates bone formation and regeneration by promoting SPP1/integrin β1-mediated osteogenic differentiation^52^. Importantly, SPP1 as a negatively charged matrix glycoprotein, has a high affinity to calcium which allows it to anchor to and retain in bone environment through the circulation system^40^. On the other hand, systemically administered LEPTIN in rodents and primates has also been reported to be localized and retained in the bone marrow^53^. Hence, LEPTIN and SPP1 may prefer to exert functional roles in the bone microenvironment. Thus, adipocyte expansion is detrimental to bone and vasculature, and further exacerbates BMAds accumulation at the expense of osteoblasts, forming a vicious feedforward loop for bone deterioration. Targeting adipocyte ESRRA might impede the disease-promoting alterations in the adipocyte-rich bone microenvironment via concomitantly modulating two pathways LEPTIN and SPP1.

## Methods

### Adipocyte-specific *Esrra* knockout mice generation

Homozygous *Esrra*^flox/flox^ mice were described previously ^23^. Both ends of the exon 2 of *Esrra* gene are flanked by loxP sites that are oriented in the same direction. The Adipoq-Cre mice [B6; FVB-Tg(Adipoq-cre)1Evdr/J] (JAX Strain: 010803) were obtained from the National Resource Center for Mutant Mice of China. The Cre/loxP system was used to generate adipocyte-specific *Esrra* knockout mice (Adipoq-Cre; *Esrra*^fl/fl^, refer as *Esrra*^AKO^) by consecutive mating of *Esrra*^flox/flox^ mice with AdipoqCre mice. The Cre-negative *Esrra*^flox/flox^ (*Esrra*^fl/fl^) mice were employed as the control genotype. The mice were housed with food and water available ad libitum and no more than 5 in a cage. All procedures used for animals and their care in this study were reviewed and approved by the ethical committee at Shenzhen Institute of Advanced Technology, Chinese Academy of Sciences.

### Animal models and treatments

For DIO, 9-week-old male *Esrra*^fl/fl^ and *Esrra*^AKO^ mice were fed a HFD (60% kcal from fat, Research Diets, Research Diet #D12492) for up to 16 weeks. Twelve weeks after HFD feeding, an oral glucose tolerance test was performed on mice fasted overnight. Blood glucose levels were measured from the tail vein before and at 30, 60, 90 and 120 min after administering glucose (2g/kg body weight), using the ACCU-CHEK blood glucose meter (ROCHE #06870279001). For OVX, 10-week-old female *Esrra*^fl/fl^ and *Esrra*^AKO^ mice were subjected to either bilateral ovariectomy to mimic postmenopausal osteoporosis or sham operation. Eight weeks post-operation, the mice were euthanized.

For pharmacological experiment, wild-type C57BL/6 mice were purchased from GemPharmatech (Nanjing, China). 7-week-old male mice were randomized into a control group and DIO group. The control mice were fed a regular chow diet throughout the experiment. The DIO mice were fed a high-fat diet for 14 weeks to induce obesity, and were then divided into two subgroups: the DIO+C29 group and the DIO+Veh group. The mice in DIO+C29 group were administered an ESRRA inverse agonist C29 (ChemPartner #S1039) through oral gavage at a dose of 30 mg/kg body weight daily for 4 weeks. The DIO+Veh group was administered an equal volume of vehicle, which was composed of 10% vitamin E-TPGS, 20% PEG400, and 70% water. Both DIO+Veh group and DIO+C29 group were exposed to 4 additional weeks of HFD feeding and treatments. All mice were fasted overnight before being sacrificed and then samples were collected for analysis. Femurs were collected for micro-CT, histological and immunofluorescence analysis. Plasma glucose, triglyceride (TG, Applygen #E1003) or free fatty acid (FFA, WAKO #294-63601) were measured using commercial kits.

### Bone micro-CT analysis

Femurs were fixed in 4% paraformaldehyde for 48 hours. Micro-CT analysis of fixed femur was performed using a Skyscan 1176 (Bruker, Kontich, Belgium) at 50 kVp, 450 μA and a resolution of 9 μm. The region of interest (ROI) was then segmented and analyzed using both the DataViewer software and CTAn software. For assessing trabecular bone, the ROI starting point was defined as 0.36 mm from the metaphyseal growth plate and extended 1 mm distal from that start point. Regions 0.5 mm above and 0.5 mm below the mid-diaphysis (50% of length) were selected to evaluate cortical bone. Three-dimensional model visualization software was applied to display the metaphyseal trabecular bone and the diaphyseal cortical bone.

### Bone histology and histomorphometry

Fixed femurs were decalcified for 3-4 weeks at 4℃ using 0.5 M EDTA (pH 7.2) before being sectioned at 5 μm. The paraffin-embedded femurs were subjected to deparaffinization and rehydration procedures before being stained using hematoxylin and eosin (H&E) kit (Beyotime #C0107) or tartrate-resistant acid phosphatase (TRAP) staining kit (Solarbio #G1492). Images were obtained using a fluorescent microscope (Olympus #BX53). The quantification of osteoblasts or osteoclasts was then measured and calculated using ImageJ software. For calcein double labeling, 20 mg/kg of calcein (Sigma-Aldrich #C0875) dissolved in 2% sodium bicarbonate solution, was administered to the mice via intraperitoneal injection at 8 and 3 days before euthanasia. Femurs were soaked in 70% ethanol, directly embedded in methyl methacrylate, and sagittally sectioned at 10 μm. Images of calcein double labeling in undecalcified femora sections were acquired on a fluorescent microscope. Dynamic indices of overall bone formation rate (BFR) and mineral apposition rate (MAR) were calibrated by ImageJ software.

### Bone immunofluorescence

For immunofluorescence staining, 5-μm bone sections were soaked in sodium citrate buffer at 65°C overnight for antigen retrieval. After being permeabilized with 0.2% Triton X-100 and blocked with 5% BSA for 1 hour, the sections were then incubated overnight at 4 °C with primary antibodies SPP1 (1:200; Novus Biologicals #AF808), PLIN1 (1: 200; Cell Signaling Technology #9349), OSTERIX (1:200; Abcam #ab209484), Endomucin (V.7C7) (EMCN,1:50; Santa Cruze #sc-65495), or CD31/PECAM-1 Alexa Fluor 488-conjugated Antibody (1:200; R&D system #FAB3628G). After primary antibody incubation, sections were washed and incubated with appropriate secondary antibodies Alexa Fluor 488-conjugated donkey anti-goat IgG (1:500; ThermoFisher #A-11055), Alexa Fluor 555-conjugated donkey anti-rabbit IgG (1:500; Abcam #ab150074), or Alexa Fluor 555-conjugated goat anti-rat IgG (1:500; Bioss #bs-0293G-AF555) for 1 hour at room temperature. Sections were then washed and nuclei were counterstained with DAPI (Cell Signaling Technology #4083) for 30 min at room temperature. After counterstaining, sections were washed and mounted with anti-fade reagent (Invitrogen #P10144). The images were captured by a fluorescent microscope.

The quantification of type H vessels was performed using ImageJ software. The area around the vessels with a range of 20 mm was lined out and the fluorescence signals based on color recognition within this area were quantified using the “area sum” parameter. The number and area of PLIN1 positive adipocytes were counted manually in each bone marrow section of distal femur by using ImageJ. For some indexes, after the signals were absolutely quantified, relative quantification was performed by taking the numerical value of one group as the reference and calculating the fold changes of other groups.

### White adipose tissue histology and immunofluorescence

gWAT was fixed in 4% paraformaldehyde for 24 hours. H&E staining was performed on paraffin-embedded gWAT sections, and the sections were also subjected to immunofluorescent staining using primary antibodies against SPP1 and LEPTIN (1:200; Abcam #ab16227). Adipose cell size and fluorescence intensity were measured using ImageJ software in a semi-automated manner.

### BMSCs isolation and culture

Mouse bone marrow MSCs (BMSCs) were isolated from tibia and femur. Briefly, cells were rinsed out with sterile syringe and then cultured in α-MEM (Gibco #C12571500BT) with 10% FBS (Gibco #10099141C) and 1% penicillin/streptomycin (Hyclone # SV30010). Cells were incubated in 5% CO_2_ incubators at 37℃. After 8 hours, non-adherent cells in the supernatants were transferred to a new culture dish, and adherent cells were further cultured with fresh medium. Following overnight culture, all non-adherent cells were washed out, and adherent cells were then fed with fresh medium. At 80%-90% confluence, the BMSCs were digested with 0.25% Trypsin (Invitrogen #25200056) and passaged with medium being changed every 48-72 hours. Human BMSCs (hBMSCs) were purchased from Cyagen Biosciences Inc. (#HUXMF-01001) and were observed to have a high level of purity, with >70% expressing CD29, CD44, and CD105, and <5% expressing CD34 and CD45 in flow cytometry assays. The maintaining of hBMSCs in growth medium was carried out with the procedure described above. Cells within 3 passages were used for *in vitro* experiments.

### Adipogenic and osteogenic differentiation

For osteogenic differentiation, BMSCs were cultured in osteogenic induction medium comprised of dexamethasone (100 nM), β-glycerophosphate (10 mM, Sigma # G9422) and ascorbic acid (50 μg/mL, Sigma #A8960). The medium was changed every 3 days for 14 days. Differentiated osteoblasts were stained with 2% alizarin red S (Beyotime #C0138). The amount of mineral content was measured by eluting the alizarin red S with 10% cetylpyridinium chloride and the optical density was measured at OD 550nm. For adipogenic differentiation of BMSCs, cells were cultured in adipogenic induction medium containing insulin (10 mg/mL, Yeasen #40112ES25), IBMX (0.5 mM, Sigma #I5879), dexamethasone (1 μM, Sigma # D4902), in the presence or absence of 1 μM rosiglitazone (Rosi, Sigma# R2408) for 7 or 14 days according to experimental design. For adipogenic differentiation of 3T3-L1 cells, cells were stimulated with adipogenic induction medium for 14 days. When specified, matured 3T3-L1 adipocytes or BMAds were treated with C29 or vehicle for an additional 2 days. Alternatively, they were infected with adenovirus expressing *Esrra* (50 pfu) or *Gfp* for 48 hours. Adenoviruses were produced and purified as we described previously^23^. Adipogenic differentiation was assessed by oil red O staining following the manufacturer’s instructions (Solarbio #G1262).

### Conditioned medium preparation

To prepare gWAT conditioned medium (CM), gWAT from *Esrra*^fl/fl^-OVX and *Esrra*^AKO^-OVX mice were minced and then incubated in α-MEM for 48 hours before collection. To prepare BMAds-CM, BMSCs were isolated from *Esrra*^fl/fl^ and *Esrra*^AKO^ mice and then subjected to adipogenesis for 14 days to achieve fully differentiated BMAds. The adipogenic induction medium was replaced with fresh α-MEM for an additional 2 days before collection. The collected CM were then centrifuged for 5 minutes at 1000 × g to remove cell pellets. The resulting supernatant was then stored at -80℃.

### Mixed differentiation

Before preparing the mixed differentiation medium, gWAT-CM was diluted 10-fold with fresh medium to obtain 10% gWAT-CM. The mixed induction medium was composed of 50% adipogenic medium and 50% osteogenic medium, with an adipogenic to osteogenic induction ratio of 1:1 as reported^41^. Wild-type BMSCs were differentiated in the mixed induction medium supplemented with 10% WAT-CM or BMAds-CM as indicated, in the presence of 0.5 μg/mL recombinant SPP1 protein (rSPP1, Abcam #ab281820), 1 μg/mL SPP1 neutralizing antibody (SPP1 Nab, Novus Biologicals #AF808), 0.5 μg/m recombinant LEPTIN protein (rLEPTIN, R&D Systems #490-OB-01M), 250 nM LEPTIN receptor antagonist Allo-aca (MCE #HY-P3212) or an equal volume of vehicle IgG (Cell Signaling Technology #2729). The formation of mineralized nodules was evaluated by alizarin red S staining, and adipocytes were distinguished by oil red O staining.

### Migration and tube formation assay

For transwell migration assays, gWAT-CM or BMAds-CM was added in the lower chamber supplemented with 0.5 μg/mL rSPP1, 1 μg/mL SPP1 Nab or an equal volume of vehicle IgG. Mouse bEnd.3 microvascular endothelial cells (ECs) were seeded into the upper chambers of 24-well plates containing an 8-μm membrane pore size (c) and were incubated for 24 hours. The migratory cells that had moved to the underside of the membrane were fixed and stained with 1% crystal violet (Beyotime #C0121). The number of migratory cells was determined by counting the cells that penetrated the membrane in 5 random fields per chamber using an optical microscope.

For tube formation assays, Matrigel (BD Biosciences #354230) was added to 24-well culture plates and polymerized for 1 hour at 37℃. After overnight starvation, ECs were resuspended with gWAT-CM or BMAds-CM and seeded at a density of 2 × 10^4^ cells/well onto the polymerized Matrigel in plates. Cells were then incubated with 0.5 μg/mL rSPP1 or 1 μg/mL SPP1 Nab at 37 °C for 4 hours, and then tube formation was assessed with an inverted microscope. The parameters of tube length and the number of branch points were analyzed using the angiogenesis analysis plugin of the Image J software. The number of migratory cells was determined by counting the cells that penetrated the membrane in 5 random fields per chamber.

### Enzyme-linked immunosorbent assay

ELISA was conducted to detect SPP1 (R&D systems #MOST00), LEPTIN (R&D systems #MOB00B), CTX1 (USCN #CEA665Mu) or PINP (NOVUS biologicals #NBP2-76466) from plasma or CM based on manufacturer’s instructions. The concentrations of SPP1 were measured by diluting plasma 300-fold, gWAT CM 300-fold, BMAds-CM 150-fold and matured 3T3-L1 adipocytes-CM 150-fold.

### Plasmid construction and dual luciferase reporter assay

*Esrra, Ppargc1a* and *Esr1* were cloned into pcDNA4 vector as we previously reported^23^. The predicted promoter regions of *Leptin* and *Spp1* were cloned into pGL3-basic luciferase reporter vector, generating pGL3-*Leptin* WT-luc and pGL3-*Spp1* WT-luc. Truncated constructs of *leptin* promoter-luciferase reporter were generated using the indicated primers listed in Table S1. Serial constructs include a promoter fragment containing site 1-3 (pGL3-Leptin S123-luc), a promoter fragment containing site 1-2 (pGL3-Leptin S12-luc), and a promoter fragment containing site 1 (pGL3-Leptin S1-luc). The putative ESRRA binding site 1 was mutated using PCR-based site-directed mutagenesis and QuikChange Site-Directed Mutagenesis Kit (Stratagene #200518) to obtain mutated constructs pGL3-*Leptin* mut 1-luc.

For transfection, 3T3-L1 cells were transiently transfected with promoter-luciferase reporters and along with several expression constructs as indicated in each figure using Lipofectamine 3000 transfection reagent (Invitrogen #L3000015) and Opti-MEM reduced serum medium (Gibco #11058021). For the *Leptin* promoter reporter assay, 3T3-L1 were transfected with *Leptin* promoter reporters and either *Esrra*, *Pgargc1a* or control vector, together with C29 or DMSO for 24 hours. For the *Spp1* promoter reporter assay, the transfected cells were cultured in phenol red-free DMEM (Gibco #A1048801) supplemented with 10% charcoal-stripped FBS for 24 hours. Cells were treated with either *Esrra*, *Esr1* or control vector, supplemented with or without 17β-estradiol (E2,10 nM) for another 24 hours, and were then exposed to adipogenic medium for 48 hours. The cells were lysed for dual luciferase reporter gene assay kit (Promega #E2980), and the transfection efficiency was normalized to Renilla.

### ChIP assay

ChIP assays were performed using a ChIP assay kit (Cell Signaling Technology #9005) per the manufacturer’s instructions. Briefly, cells were fixed with 1% formaldehyde, quenched with glycine solution, and then washed with PBS. Nuclei were extracted and micrococcal nuclease digestion was performed, followed by sonication to obtain 200-500 bp genomic DNA fragments. Chromatin complexes were immunoprecipitated with primary antibody as below and incubated with protein G magnetic beads (Cell Signaling Technology #9006) overnight. To determine the repression effect of ESRRA on the *Spp1* promoter, BMSCs from *Esrra*^fl/fl^ and *Esrra*^AKO^ mice were infected with adenovirus expressing *Esrra* (50 pfu) or control *Gfp* for 24 hours and subsequently stimulated with adipogenic induction medium in phenol red-free DMEM supplemented with 10% charcoal stripped FBS with or without E2 (10 nM). After 4 days, cells were subjected to ChIP assay with primary antibody against ESR1 (2 μl/IP, Abcam #ab32063) or normal rabbit IgG (1 μg/IP, Cell Signaling Technology #2729). To determine physical binding of ESRRA on the *Lepin* promoter, 3T3-L1 cells were subjected to adipogenic induction for 4 days with treatment of 20 μM C29 or DMSO, or with addition of adenovirus expressing *Esrra* or control as described above. These cells were then subjected to ChIP assays with primary antibody against ESRRA (10 μl/IP, Cell Signaling Technology #13826) or normal rabbit IgG. The precipitated DNA was purified and used as a template for PCR using primers specifically designed to amplify a segment of 150-250 bp covering the putative ESRRA or ESR1 binding sites. Primer sequences used for ChIP-qPCR are listed in Table S2. The amount of immunoprecipitated DNA in each sample was normalized to IgG and presented as fold relative enrichment.

### RNA isolation and qRT-PCR

Total RNA was extracted from homogenized gWAT or cells using AG RNAex Pro Reagent (Accurate Biotechnology #AG21101). Reverse transcription was performed from 2 μg of total RNAs using HiScript III 1st Strand cDNA Synthesis Kit (Vazyme #R312). qRT-PCR was employed to quantify the mRNA levels. RealStar Green Fast Mixture (Genstar #A301-10) was used as the detection reagent, and the primers listed in table S3 were applied. The quantity of mRNA was calculated using the 2^−ΔΔCt method. All reactions were performed at least four independent experiments.

### RNA-seq and gene set enrichment analysis

Total RNA was extracted using RNA isolation kit (Vazyme #RC112) from BMSCs from *Esrra*^fl/fl^ and *Esrra*^AKO^ mice which were subjected to adipogenic induction for 7 days. The total RNA quantity and purity were analyzed by using Bioanalyzer 2100 and RNA 6000 Nano LabChip Kit (Agilent, CA, USA, 5067-1511). RNA-seq libraries were constructed and then sequenced using an Illumina Novaseq™ 6000 (LC-Bio Technology, China) following the vendor’s recommended protocol. FPKM (fragments per million reads per kilobase mapped) method was used to normalize gene expression, and lowly expressed genes (average FPKM < 10) were filtered in each sample. Then difference expression of genes (DEG) were analyzed using DESeq2 software and then subjected to enrichment analysis of GO functions and KEGG pathways. Significantly enriched GO terms and KEGG pathways were selected as follows: *P* value < 0.05. A heatmap analysis was performed to visualize the Z-score calculation values for gene expression of secreted factors, which were identified across two experimental group comparisons (*P* value < 0.05, fold change > 1.5).

### Immunoblot analysis

Cells or tissues were collected and lysed in RIPA buffer (Beyotime #P0013B) containing a cocktail of protease inhibitors and phosphatase inhibitors. The protein concentration was quantified by BCA Assay (Thermo Fisher #23227). The lysate was mixed with SDS loading buffer and heated at 100℃ for 10 minutes. Subsequently, 30-60 μg of proteins were loaded onto gel for SDS-PAGE analysis. After SDS-PAGE analysis, proteins were transferred onto a PVDF membrane (Merck Millipore #IPFL00010) and blocked with 5% skimmed milk-PBST. Then, the membrane was incubated overnight with primary antibodies that were dissolved in 5% BSA. The antibodies used for immunoblot analysis were anti-ESRRA (1:1000), anti-TUBULIN (1:5000, BPI #AbM9005-37B-PU), anti-LEPTIN (1:500), anti-SPP1 (1:1000) or anti-GAPDH (1:5000, Proteintech #60004-1-Ig). The immunocomplexes were incubated with corresponding secondary antibodies: HRP-goat anti-mouse IgG (EarthOx Life #E030110-02), HRP-goat anti-rabbit IgG (EarthOx Life #E030120-02) or HRP-rabbit anti-goat IgG (ABclonal #AS029). They were then detected with ECL luminescence reagent (Millipore #WBLUR0500). Images of the samples were captured using ChemiDoc XRS chemiluminescence imaging system (Bio-Rad).

### Statistical analysis

Differences between groups were assessed by unpaired Student’s t-test using Graph Prism 6. For the data from multigroup studies, one-way ANOVA analysis was followed by Bonferroni post hoc analysis. *P* value <0.05 was considered statistically significant.

## Acknowledgements

This work was supported in part by grants from the National Key Research and Development Program of China (2018YFA0703100), National Natural Science Foundation of China (82072493, 81770882, 81570532 and 81972071), Shenzhen Science and Technology Program (JCYJ20210324101800002), Guangdong Basic and Applied Basic Research Foundation (2022A1515010528), Fund of State Key Laboratory of Phytochemistry and Plant Resources in West China (P2022-KF12).

## Author contributions

T.H. and M.G. conceived, designed the work, analyzed results, and wrote the manuscript. T.H. carried out most of the experiments. Z.L, Z.W., L.G. and J.G. helped with some experiments. N.Z. and L.C. helped with bioinformatics analysis. C.W. proofread the manuscript. M.G., X.Z., H.W, C.G., H.P., K.W.K.Y. and W.L. secured funding. All authors reviewed the manuscript.

## References

1. Tuckermann J, Adams RH. The endothelium-bone axis in development, homeostasis and bone and joint disease. Nature reviews Rheumatology 2021, 17(10): 608–620.

2. Ambrosi TH, Scialdone A, Graja A, Gohlke S, Jank AM, Bocian C, et al. Adipocyte Accumulation in the Bone Marrow during Obesity and Aging Impairs Stem Cell-Based Hematopoietic and Bone Regeneration. Cell stem cell 2017, 20(6): 771–784.e776.

3. Zhou BO, Yue R, Murphy MM, Peyer JG, Morrison SJ. Leptin-receptor-expressing mesenchymal stromal cells represent the main source of bone formed by adult bone marrow. Cell stem cell 2014, 15(2): 154–168.

4. Suchacki KJ TA, Mattiucci D, Scheller EL, Papanastasiou G, Gray C, Sinton MC, Ramage LE, McDougald WA, Lovdel A, Sulston RJ, Thomas BJ, Nicholson BM, Drake AJ, Alcaide-Corral CJ, Said D, Poloni A, Cinti S, Macpherson GJ, Dweck MR, Andrews JPM, Williams MC, Wallace RJ, van Beek EJR, MacDougald OA, Morton NM, Stimson RH, Cawthorn WP. . Bone marrow adipose tissue is a unique adipose subtype with distinct roles in glucose homeostasis. Nature communications 2020, 11(1): 3097.

5. Sebo ZL, Rendina-Ruedy E, Ables GP, Lindskog DM, Rodeheffer MS, Fazeli PK, et al. Bone Marrow Adiposity: Basic and Clinical Implications. Endocrine reviews 2019, 40(5): 1187–1206.

6. Peng H, Hu B, Xie LQ, Su T, Li CJ, Liu Y, et al. A mechanosensitive lipolytic factor in the bone marrow promotes osteogenesis and lymphopoiesis. Cell metabolism 2022, 34(8): 1168–1182.e1166.

7. Yue R, Zhou BO, Shimada IS, Zhao Z, Morrison SJ. Leptin Receptor Promotes Adipogenesis and Reduces Osteogenesis by Regulating Mesenchymal Stromal Cells in Adult Bone Marrow. Cell stem cell 2016, 18(6): 782–796.

8. Cawthorn WP, Scheller EL, Learman BS, Parlee SD, Simon BR, Mori H, et al. Bone marrow adipose tissue is an endocrine organ that contributes to increased circulating adiponectin during caloric restriction. Cell metabolism 2014, 20(2): 368–375.

9. Scheller EL, Cawthorn WP, Burr AA, Horowitz MC, MacDougald OA. Marrow Adipose Tissue: Trimming the Fat. Trends in endocrinology and metabolism: TEM 2016, 27(6): 392–403.

10. Fan Y, Hanai JI, Le PT, Bi R, Maridas D, DeMambro V, et al. Parathyroid Hormone Directs Bone Marrow Mesenchymal Cell Fate. Cell metabolism 2017, 25(3): 661–672.

11. Hu Y LX, Zhi X, Cong W, Huang B, Chen H, Wang Y, Li Y, Wang L, Fang C, Guo J, Liu Y, Cui J, Cao L, Weng W, Zhou Q, Wang S, Chen X, Su J. . RANKL from bone marrow adipose lineage cells promotes osteoclast formation and bone loss. EMBO reports 2021, 22(7): e52481.

12. Kusumbe AP, Ramasamy SK, Adams RH. Coupling of angiogenesis and osteogenesis by a specific vessel subtype in bone. Nature 2014, 507(7492): 323–328.

13. Liu X, Gu Y, Kumar S, Amin S, Guo Q, Wang J, et al. Oxylipin-PPARγ-initiated adipocyte senescence propagates secondary senescence in the bone marrow. Cell metabolism 2023, 35(4): 667–684.e666.

14. Wang L, Zhou F, Zhang P, Wang H, Qu Z, Jia P, et al. Human type H vessels are a sensitive biomarker of bone mass. Cell Death and Disease 2017, 8(5): e2760.

15. Zhu Y, Ruan Z, Lin Z, Long H, Zhao R, Sun B, et al. The association between CD31(hi)Emcn(hi) endothelial cells and bone mineral density in Chinese women. Journal of bone and mineral metabolism 2019, 37(6): 987–995.

16. Naveiras O, Nardi V, Wenzel PL, Hauschka PV, Fahey F, Daley GQ. Bone-marrow adipocytes as negative regulators of the haematopoietic microenvironment. Nature 2009, 460(7252): 259–263.

17. Zhou BO, Yu H, Yue R, Zhao Z, Rios JJ, Naveiras O, et al. Bone marrow adipocytes promote the regeneration of stem cells and haematopoiesis by secreting SCF. Nature cell biology 2017, 19(8): 891–903.

18. Zhong L, Yao L, Tower RJ, Wei Y, Miao Z, Tong W, et al. Single cell transcriptomics identifies a unique adipose lineage cell population that regulates bone marrow environment. eLife 2020, 9.

19. Giguère V. Transcriptional control of energy homeostasis by the estrogen-related receptors. Endocrine reviews 2008, 29(6): 677–696.

20. Ranhotra HS. Up-regulation of orphan nuclear estrogen-related receptor alpha expression during long-term caloric restriction in mice. Molecular and cellular biochemistry 2009, 332(1-2): 59–65.

21. Luo J, Sladek R, Carrier J, Bader JA, Richard D, Giguère V. Reduced fat mass in mice lacking orphan nuclear receptor estrogen-related receptor alpha. Molecular and cellular biology 2003, 23(22): 7947–7956.

22. Emmett MJ, Lim HW, Jager J, Richter HJ, Adlanmerini M, Peed LC, et al. Histone deacetylase 3 prepares brown adipose tissue for acute thermogenic challenge. Nature 2017, 546(7659): 544–548.

23. Yang M, Liu Q, Huang T, Tan W, Qu L, Chen T, et al. Dysfunction of estrogen-related receptor alpha-dependent hepatic VLDL secretion contributes to sex disparity in NAFLD/NASH development. Theranostics 2020, 10(24): 10874–10891.

24. Kuang Z, Wang Y, Ye C, Ruhn KA, Li Y, Hooper LV, et al. The intestinal microbiota programs diurnal rhythms in host metabolism through histone deacetylase 3. Science (New York, NY) 2019, 365(6460): 1428–1434.

25. Dhillon P, Park J, Hurtado Del Pozo C, Li L, Doke T, Huang S, et al. The Nuclear Receptor ESRRA Protects from Kidney Disease by Coupling Metabolism and Differentiation. Cell metabolism 2021, 33(2): 379–394.e378.

26. Xia H, Scholtes C, Dufour CR, Ouellet C, Ghahremani M, Giguère V. Insulin action and resistance are dependent on a GSK3β-FBXW7-ERRα transcriptional axis. Nature communications 2022, 13(1): 2105.

27. Tencerova M, Figeac F, Ditzel N, Taipaleenmäki H, Nielsen TK, Kassem M. High-Fat Diet-Induced Obesity Promotes Expansion of Bone Marrow Adipose Tissue and Impairs Skeletal Stem Cell Functions in Mice. Journal of bone and mineral research : the official journal of the American Society for Bone and Mineral Research 2018, 33(6): 1154–1165.

28. Veldhuis-Vlug AG, Rosen CJ. Clinical implications of bone marrow adiposity. Journal of internal medicine 2018, 283(2): 121–139.

29. Cohen A, Dempster DW, Stein EM, Nickolas TL, Zhou H, McMahon DJ, et al. Increased marrow adiposity in premenopausal women with idiopathic osteoporosis. The Journal of clinical endocrinology and metabolism 2012, 97(8): 2782–2791.

30. Deng P, Yuan Q, Cheng Y, Li J, Liu Z, Liu Y, et al. Loss of KDM4B exacerbates bone-fat imbalance and mesenchymal stromal cell exhaustion in skeletal aging. Cell stem cell 2021, 28(6): 1057–1073.e1057.

31. Pop LM, Lingvay I, Yuan Q, Li X, Adams-Huet B, Maalouf NM. Impact of pioglitazone on bone mineral density and bone marrow fat content. Osteoporosis international : a journal established as result of cooperation between the European Foundation for Osteoporosis and the National Osteoporosis Foundation of the USA 2017, 28(11): 3261–3269.

32. Pachón-Peña G, Bredella MA. Bone marrow adipose tissue in metabolic health. Trends in endocrinology and metabolism: TEM 2022, 33(6): 401–408.

33. Chen Q, Shou P, Zhang L, Xu C, Zheng C, Han Y, et al. An osteopontin-integrin interaction plays a critical role in directing adipogenesis and osteogenesis by mesenchymal stem cells. Stem cells (Dayton, Ohio) 2014, 32(2): 327–337.

34. Deblois G, Giguère V. Oestrogen-related receptors in breast cancer: control of cellular metabolism and beyond. Nature reviews Cancer 2013, 13(1): 27–36.

35. Vanacker JM, Pettersson K, Gustafsson JA, Laudet V. Transcriptional targets shared by estrogen receptor-related receptors (ERRs) and estrogen receptor (ER) alpha, but not by ERbeta. The EMBO journal 1999, 18(15): 4270–4279.

36. Duvall CL, Taylor WR, Weiss D, Wojtowicz AM, Guldberg RE. Impaired angiogenesis, early callus formation, and late stage remodeling in fracture healing of osteopontin-deficient mice. Journal of bone and mineral research : the official journal of the American Society for Bone and Mineral Research 2007, 22(2): 286–297.

37. Hochmann S, Ou K, Wolf M, Strunk D, Schmidt-Bleek K, Schallmoser K, et al. The enhancer landscape predetermines the skeletal regeneration capacity of stromal cells. Science translational medicine 2023, 15(688): eabm7477.

38. Langen UH, Pitulescu ME, Kim JM, Enriquez-Gasca R, Sivaraj KK, Kusumbe AP, et al. Cell-matrix signals specify bone endothelial cells during developmental osteogenesis. Nature cell biology 2017, 19(3): 189–201.

39. Xie H, Cui Z, Wang L, Xia Z, Hu Y, Xian L, et al. PDGF-BB secreted by preosteoclasts induces angiogenesis during coupling with osteogenesis. Nature medicine 2014, 20(11): 1270–1278.

40. Dai B, Xu J, Li X, Huang L, Hopkins C, Wang H, et al. Macrophages in epididymal adipose tissue secrete osteopontin to regulate bone homeostasis. Nature communications 2022, 13(1): 427.

41. Yu Y, Newman H, Shen L, Sharma D, Hu G, Mirando AJ, et al. Glutamine Metabolism Regulates Proliferation and Lineage Allocation in Skeletal Stem Cells. Cell metabolism 2019, 29(4): 966-978.e964.

42. Huang T, Fu X, Wang N, Yang M, Zhang M, Wang B, et al. Andrographolide prevents bone loss via targeting estrogen-related receptor-α-regulated metabolic adaption of osteoclastogenesis. British journal of pharmacology 2021, 178(21): 4352–4367.

43. Bae S, Lee MJ, Mun SH, Giannopoulou EG, Yong-Gonzalez V, Cross JR, et al. MYC-dependent oxidative metabolism regulates osteoclastogenesis via nuclear receptor ERRα. The Journal of clinical investigation 2017, 127(7): 2555–2568.

44. Scheller EL, Rosen CJ. What’s the matter with MAT? Marrow adipose tissue, metabolism, and skeletal health. Annals of the New York Academy of Sciences 2014, 1311(1): 14–30.

45. Shen W, Chen J, Gantz M, Punyanitya M, Heymsfield SB, Gallagher D, et al. MRI-measured pelvic bone marrow adipose tissue is inversely related to DXA-measured bone mineral in younger and older adults. European journal of clinical nutrition 2012, 66(9): 983–988.

46. Laharrague P, Larrouy D, Fontanilles AM, Truel N, Campfield A, Tenenbaum R, et al. High expression of leptin by human bone marrow adipocytes in primary culture. FASEB journal : official publication of the Federation of American Societies for Experimental Biology 1998, 12(9): 747–752.

47. Miggitsch C, Meryk A, Naismith E, Pangrazzi L, Ejaz A, Jenewein B, et al. Human bone marrow adipocytes display distinct immune regulatory properties. EBioMedicine 2019, 46: 387–398.

48. Tencerova M, Frost M, Figeac F, Nielsen TK, Ali D, Lauterlein JL, et al. Obesity-Associated Hypermetabolism and Accelerated Senescence of Bone Marrow Stromal Stem Cells Suggest a Potential Mechanism for Bone Fragility. Cell reports 2019, 27(7): 2050–2062.e2056.

49. Yuan S, Stewart KS, Yang Y, Abdusselamoglu MD, Parigi SM, Feinberg TY, et al. Ras drives malignancy through stem cell crosstalk with the microenvironment. Nature 2022, 612(7940): 555–563.

50. Kratofil RM, Shim HB, Shim R, Lee WY, Labit E, Sinha S, et al. A monocyte-leptin-angiogenesis pathway critical for repair post-infection. Nature 2022, 609(7925): 166–173.

51. Villena JA, Kralli A. ERRalpha: a metabolic function for the oldest orphan. Trends in endocrinology and metabolism: TEM 2008, 19(8): 269–276.

52. Chen L, Shi K, Ditzel N, Qiu W, Figeac F, Nielsen LHD, et al. KIAA1199 deficiency enhances skeletal stem cell differentiation to osteoblasts and promotes bone regeneration. Nature communications 2023, 14(1): 2016.

53. Ceccarini G, Flavell RR, Butelman ER, Synan M, Willnow TE, Bar-Dagan M, et al. PET imaging of leptin biodistribution and metabolism in rodents and primates. Cell metabolism 2009, 10(2): 148–159.

