## Supplementary Information for "Targeting adipocyte ESRRA promotes osteogenesis and vascular formation in adipocyte-rich bone marrow"

**Supplementary information contents:**

**1. Supplementary Figures: 1-10**
**2. Supplementary Tables: 1-3**

**Fig. S1**

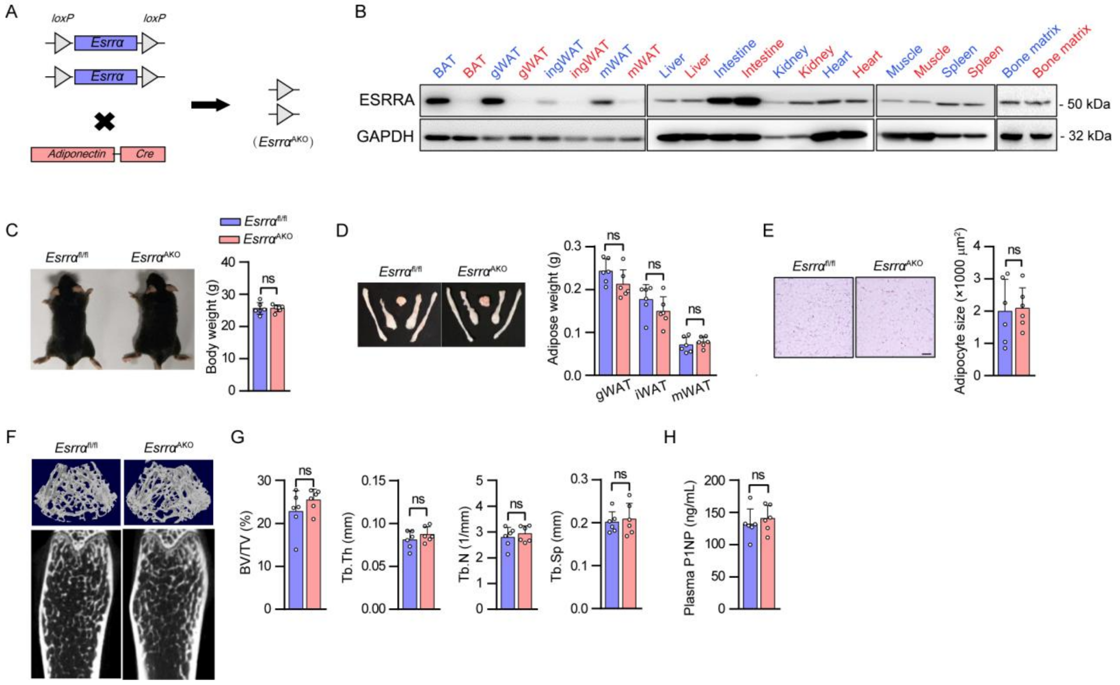


**Fig. S1. *Esrra*$^{AKO}$ male mice exhibit normal WAT and bone phenotype at the age of 10-week-old.** (A) Schematic diagram showing the breeding strategy to generate adipocyte-specific knockout *Esrra* mice. *Esrra*$^{flox/flox}$ mice were continuously mated with AdipoqCre mice to obtain *Esrra*$^{flox/flox}$; AdipoCre mice (*Esrra*$^{AKO}$) and littermate control *Esrra*$^{flox/flox}$ mice (*Esrra*$^{fl/fl}$). Phenotypic analysis was conducted on 10-week-old mice. (B) The protein expression levels of ESRRA were evaluated in adipose tissues and non-adipose tissues, comparing *Esrra*$^{fl/fl}$ mice (blue font) with *Esrra*$^{AKO}$ mice (red font). (C) Representative images and quantitative analysis of body weights in *Esrra*$^{fl/fl}$ and *Esrra*$^{AKO}$ male mice. (D) Representative images of white adipose tissue depots. The weights of gWAT, iWAT and mWAT are shown on the right panel. (E) H&E staining of gWAT sections and quantitative analysis of adipocyte size (scale bar: 50 μm). (F) Representative single micro-CT sagittal section and 3-dimensional reconstitution of distal femurs. (G) Quantitative micro-CT analysis of BV/TV, Tb.Th, Tb.N and Tb.Sp. (H) Plasma P1NP levels. Data are shown as mean ± SD (n = 6). *$P$ <0.05 was considered statistically significant.

**Fig. S2**

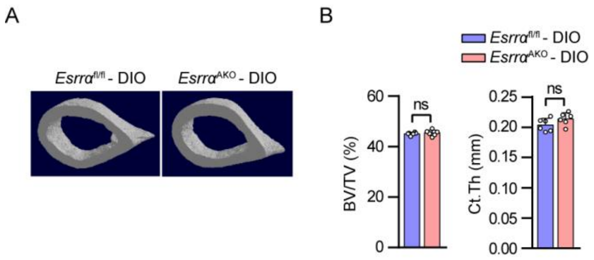


**Fig. S2. Adipocyte ESRRA ablation results in no alterations in cortical bone in DIO mice.** (A) Representative μCT images of cortical bone in femurs from *Esrra*$^{fl/fl}$ DIO and *Esrra*$^{AKO}$ DIO mice. (B) Bone histomorphometric analysis of BV/TV and Ct.Th of cortical bone in femoral midshaft. Data are shown as mean ± SD (n = 6). **P* <0.05 was considered statistically significant.

**Fig. S3**

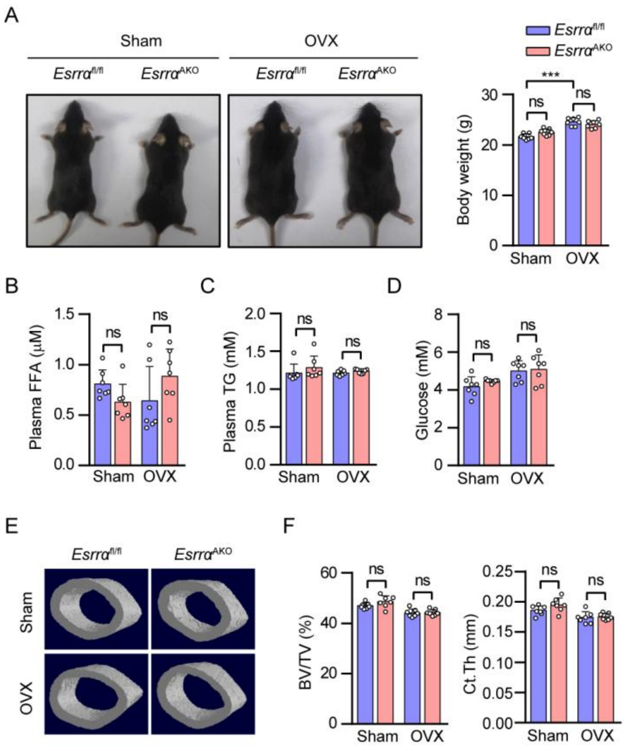


**Fig. S3. *Esrra*$^{AKO}$ female mice display no significant changes in blood biochemistry and cortical bone compared to *Esrra*$^{fl/fl}$ mice following OVX.** (A) Representative photographs and body weights analysis of *Esrra*$^{fl/fl}$ and *Esrra*$^{AKO}$ female mice underwent either sham or OVX operation for 8 weeks. (B-C) Plasma FFA (B) and TG (C) levels. (D) Blood glucose levels. (E-F) Representative μCT images (E) and bone histomorphometric analysis of BV/TV and Ct.Th (F) of cortical bone in femoral midshaft. Data are shown as mean ± SD (n = 7 per group). *$P$ <0.05 was considered statistically significant.

**Fig. S4**

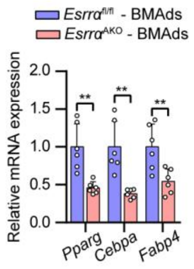


**Fig. S4. The expression levels of adipogenic markers in BMAds from *Esrra*$^{AKO}$ and *Esrra*$^{fl/fl}$ mice.** mRNA expression levels of adipogenic markers *Pparg, Cebpa* and *Fabp4* were measured in BMAds from *Esrra*$^{fl/fl}$ and *Esrra*$^{AKO}$ mice (n = 6). Data are shown as mean ± SD. *$P$ <0.05 was considered statistically significant.

**Fig. S5**

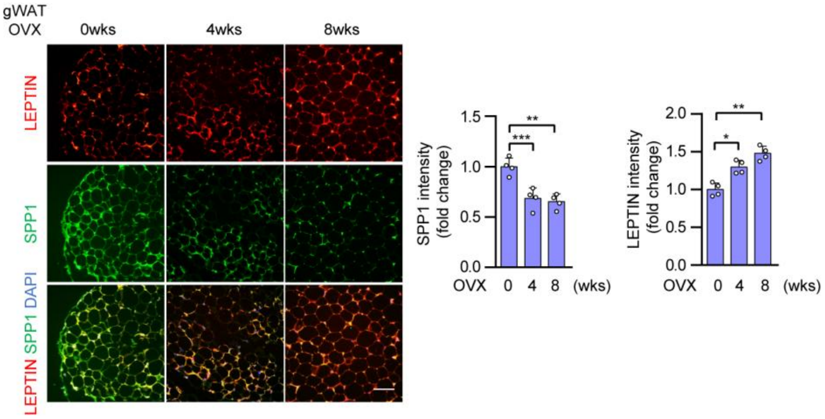


**Fig. S5. Immunofluorescence analysis of SPP1 and LEPTIN in gWAT of wild-type female mice following OVX.** Immunofluorescence co-staining and quantification of SPP1 (green) and LEPTIN (red) in gWAT sections of wild-type mice at 0 week, 4 weeks, and 8 weeks after OVX surgery. Scale bar: 50 µm. Data are shown as mean ± SD (n = 4 mice). *$P$ <0.05 was considered statistically significant.

**Fig. S6**

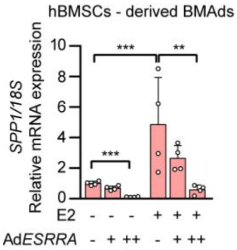


**Fig. S6. *SPP1* expression is repressed by ESRRA in human BMSCs-derived BMAds.** mRNA expression levels of *Spp1* were measured in human BMSCs-derived BMAds infected with adenovirus expressing *ESRRA* or control *GFP* in the presence of the indicated E2 treatments for 2 days. n = 4 biologically independent samples. Data are shown as mean ± SD. **P* <0.05 was considered statistically significant.

**Fig. S7**

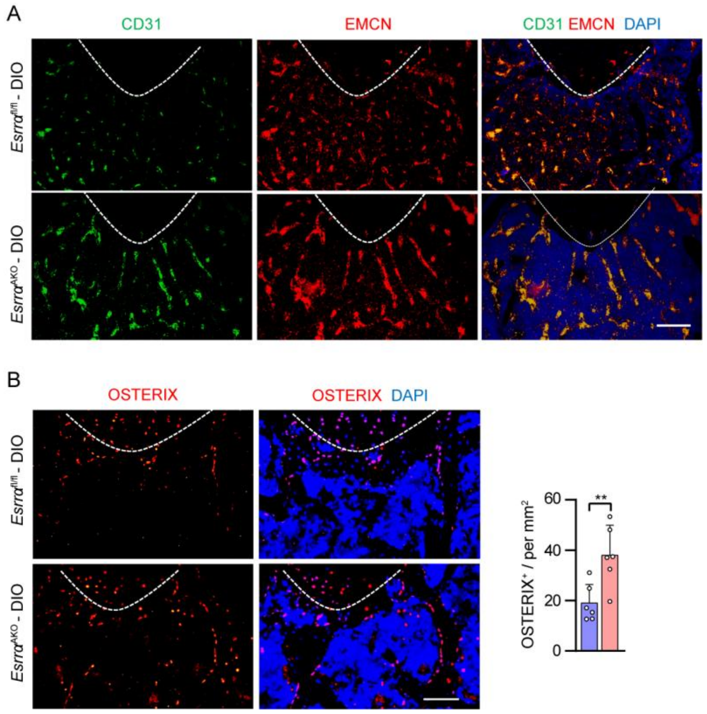


**Fig. S7. Immunofluorescence staining for type H vessel and osteoprogenitors in distal femurs of *Esrra*$^{AKO}$ DIO mice and corresponding control mice.** (A) Representative images of metaphyseal type H vessels immunostained for EMCN (red) and CD31 (green) in distal femurs of *Esrra*$^{fl/fl}$-DIO and *Esrra*$^{AKO}$-DIO mice. DAPI (blue) is used for counterstaining of nuclei. Scale bar: 100 μm. (B) Immunostaining and quantitation analysis of OSTERIX (red) with DAPI (blue) in the metaphysis of distal femurs. Scale bar: 50 μm. Data are shown as mean ± SD (n=6 per group). $^{*}P$ <0.05 was considered statistically significant.

**Fig. S8**

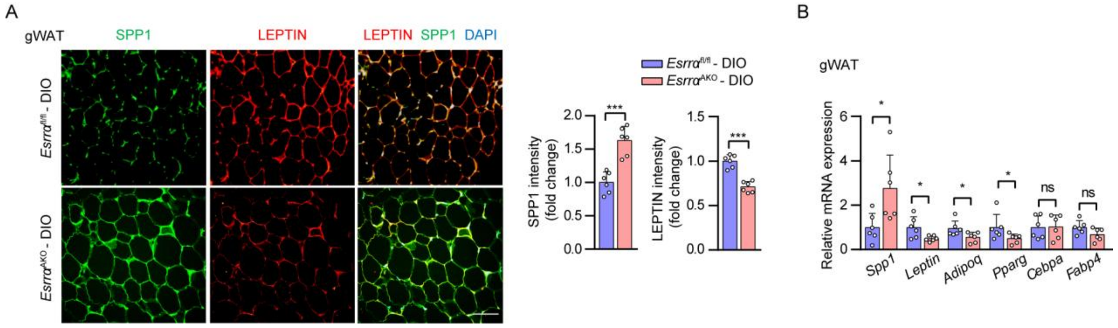


**Fig. S8. Immunofluorescence and mRNA levels analysis in gWAT of *Esrra*$^{AKO}$ DIO mice and corresponding control mice.** (A) Immunofluorescence co-staining of SPP1 and LEPTIN with quantitative analysis of fluorescence intensity in gWAT of *Esrra*$^{fl/fl}$ - DIO and *Esrra*$^{AKO}$ - DIO mice (scale bar: 100 μm). (B) mRNA expression of *Spp1, Leptin, Pparg, Cebpa* and *Fabp4* were examined in gWAT. Data are shown as mean ± SD (n=6 per group). **P* <0.05 was considered statistically significant.

**Fig. S9**

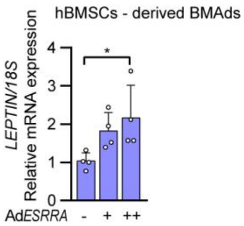


**Fig. S9. mRNA expression of *LEPTIN* is induced by ESRRA overexpressing in human BMSCs-derived BMAds.** mRNA expression levels of *LEPTIN* were measured in matured human BMSCs-derived BMAds infected with adenovirus expressing *ESRRA* or control *GFP* for 2 days. n = 4. Data are shown as mean ± SD. **P* <0.05 was considered statistically significant.

**Fig. S10**

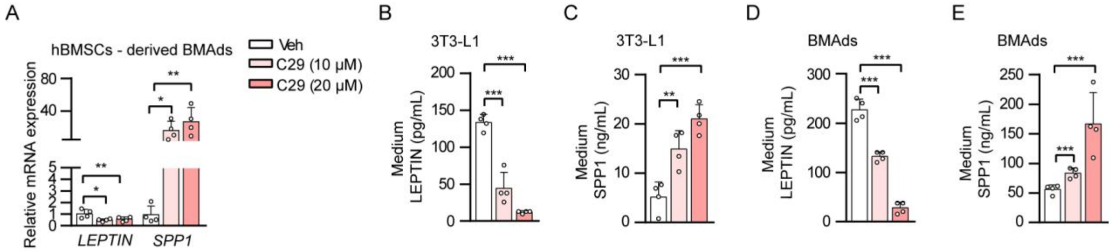


**Fig. S10. mRNA expression and secretion levels of SPP1 and LEPTIN in adipocytes treated with C29.** (A) mRNA levels of *LEPTIN* and *SPP1* were measured in human BMSCs-derived BMAds treated with C29 or DMSO for 2 days. n = 4. (B-C) The concentrations of soluble SPP1(B) and LEPTIN (C) in the culture medium of mature 3T3-L1 adipocytes were measured by ELISA (n=4). (D-E) The concentrations of soluble SPP1 (D) and LEPTIN (E) in BMAds-CM (n=4). Data are shown as mean ± SD. *$P$ <0.05 was considered statistically significant.

### Supplementary Table 1. Primer sequences of promoter vectors

| Primer name | Sequence 5'- 3' |
|---|---|
| *Spp1* WT-luc | F: CGGGGTACCGGGGTCATATGGTTCAGCTC |
| | R: CCCAAGCTTAGACTGCAAACCCAAGCAAG |
| *Leptin* WT-luc | F: CTAGCTAGCTGGGCATGATGCGACCATTT |
| | R: CCCAAGCTTAGCTGCTGGAGCAGGGA |
| *Leptin* S123-luc | F: CTAGCTAGCCTTCGGGTACCAAAGGAAGACA |
| | R: CCCAAGCTTAGCTGCTGGAGCAGGGAT |
| *Leptin* S12-luc | F: CTAGCTAGCCCTCTGAGCAGCCAGGTTAGG |
| | R: CCCAAGCTTAGCTGCTGGAGCAGGGAT |
| *Leptin* S1-luc | F: CTAGCTAGCGCAAAGAGCTGTCGGAAAAA |
| | R: CCCAAGCTTAGCTGCTGGAGCAGGGAT |
| *Leptin* mutS1-luc | F: GCTGCTGGCCGGAAATCGAGGATTACCGG |
| | R: CCGGTAATCCTCGATTTCCGGCCAGCAGC |

**Supplementary Table 2.    Primer sequences of ChIP-qPCR**

| Primer name | Sequence 5’- 3’ |
|---|---|
| *Leptin* R1 | F: TGGCCGGACCTCGAGGATTA |
| | R: CTTGCGCAACTGTCCGGC |
| *Leptin* R2 | F: GCAGGTGCATTCTGTGATGTC |
| | R: GCTCTTTGCATACCTAACCTGG |
| *Leptin* R3 | F: gTTTCCTCCCATTAggAACCCA |
| | R: CgAAGGTCGCAAGTGTGTTT |
| *Spp1* R1 | F: CCAACTGACCTGGAACACAGT |
| | R: GTGGCTCTGTTTTGTACTCCG |
| *Spp1* R2 | F: AGCAACAAGGTTCACGAGGT |
| | R: TATGCAGCCGCTTGCTCTTT |

### Supplementary Table 3. Primer sequences for qRT-PCR

| Primer name | Sequence 5'-3' |
|---|---|
| Mouse *18s* | F: TAAGTCCCTGCCCTTTGTACACA |
| | R: GATCCGAGGGCCTCACTAAAC |
| Mouse *Esrra* | F: CTCAGCTCTCTACCCAAACGC |
| | R: CCGCTTGGTGATCTCACACTC |
| Mouse *Adipoq* | F: TGTTCCTCTTAATCCTGCCCA |
| | R: CCAACCTGCACAAGTTCCCTT |
| Mouse *Leptin* | F: GAGACCCCTGTGTCGGTTC |
| | R: CTGCGTGTGTGAAATGTCATTG |
| Mouse *Spp1* | F: AGCAAGAAACTCTTCCAAGCAA |
| | R: GTGAGATTCGTCAGATTCATCCG |
| Mouse *Pparg* | F: TCGCTGATGCACTGCCTATG |
| | R: GAGAGGTCCACAGAGCTGATT |
| Mouse *CEBPa* | F: CCGTGGTGGTTTCTCCTTGA |
| | R: TCATTTTTCTCTCACGGGGCCA |
| Mouse *Fabp4* | F: TGAAATCACCGCAGACGACA |
| | R: ACACATTCCACCACCAGCTT |
| Mouse *Sp7* | F: ATGGCGTCCTCTCTGCTTG |
| | R: TGAAAGGTCAGCGTATGGCTT |
| Mouse *Bglap* | F: CCATCTTTCTGCTCACTCT |
| | R: GTCTGTTCACTACCTTATTGC |
| Mouse *Runx2* | F: AGAGTCAGATTACAGATCCCAGG |
| | R: TGGCTCTTCTTACTGAGAGAGG |
| Human *SPP1* | F: GAAGTTTCGCAGACCTGACAT |
| | R: GTATGCACCATTCAACTCCTCG |
| Human *LEPTIN* | F: TGCCTTCCAGAAACGTGATCC |
| | R: CTCTGTGGAGTAGCCTGAAG |
